# Targeting non-canonical NF-κB signalling in CYLD cutaneous syndrome by selective inhibition of IκB kinase alpha

**DOI:** 10.1101/2025.01.31.635629

**Authors:** Kirsty Hodgson, Joseph Inns, Gary Reynolds, Emily Stephenson, Andrew Paul, Naomi Sinclair, Giacomo Berretta, Christopher Lawson, Andrew Michael Frey, Iglika Ivanova, Eva Adam, Christopher J. Lord, Simon Cockell, Jonathan Coxhead, Nikoletta Nagy, David Adams, Marta Szell, Matthias Trost, Muzlifah Haniffa, Simon P. Mackay, Neil Perkins, Neil Rajan

## Abstract

CYLD cutaneous syndrome (CCS) skin tumors develop from puberty onwards, can number in the hundreds and progressively grow over time. CCS patients lack medical therapies and require repeated surgery to control tumor burden. *CYLD* loss of heterozygosity (LOH) drives tumor growth, and CCS tumors have previously been shown to demonstrate increased canonical NF-κB and Wnt signalling. Here, we demonstrate evidence of non-canonical NF-κB signalling in CCS tumor keratinocytes, with increased p100 to p52 processing and RelB protein expression compared to normal skin. Utilizing complementary transcriptomics and proteomics on patient derived CCS tumor cell fractions, we identify IκB kinase alpha (IKKα) as a candidate target in the non-canonical NF-κB signalling pathway. A novel, highly selective, IKKα inhibitor (SU1644) used in patient derived CCS tumor spheroid cultures demonstrated that IKKα inhibition reduced tumor spheroid viability. These data provide the pre-clinical rationale for the assessment of topical IKKα inhibitors as a novel preventative treatment for CCS.

**Teaser:** Topical IKKα inhibition emerges as a potential therapy for CYLD cutaneous syndrome by targeting non-canonical NF-κB signalling

**Graphical abstract:** 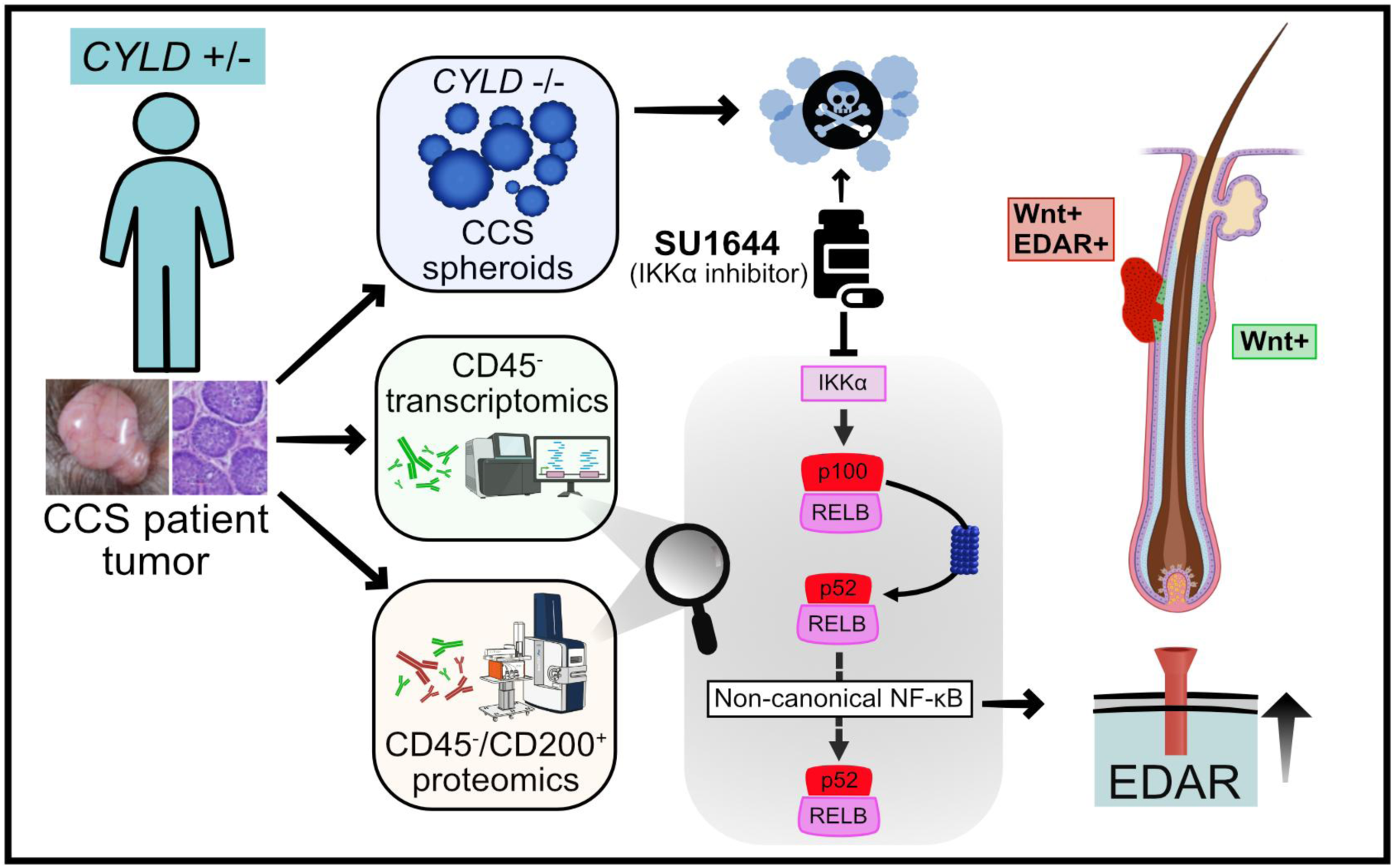

## Introduction

CYLD cutaneous syndrome (CCS) is a dominantly inherited skin tumor syndrome driven primarily by loss of function variants (PVs) in *CYLD* (**Figure 1A**). *CYLD* encodes a deubiquitinase (DUB) that negatively regulates NF-κB signalling (1). The majority of CCS patients carry germline heterozygous truncating *CYLD* PVs involving the DUB domain, and CCS tumors result from loss of heterozygosity (LOH), resulting in CYLD inactivation (2). Patients progressively develop multiple hair follicle tumors (cylindromas and spiradenomas with extensive CD200+ expression (3) on the head and torso (4), which enlarge from puberty onwards (5). Benign CCS skin tumors can transform, with malignant CCS tumors invading the skull, metastasizing to the lung (6), and other sites resulting in death. CCS patients have stated the significant impact on quality of life of repeated lifelong surgeries that may culminate in total scalp removal is poorly understood by the medical profession (7). There are no medical therapies that can be given pre-emptively to known *CYLD* PV carriers, and CCS represents an area of unmet medical need (8).

**Figure 1.**
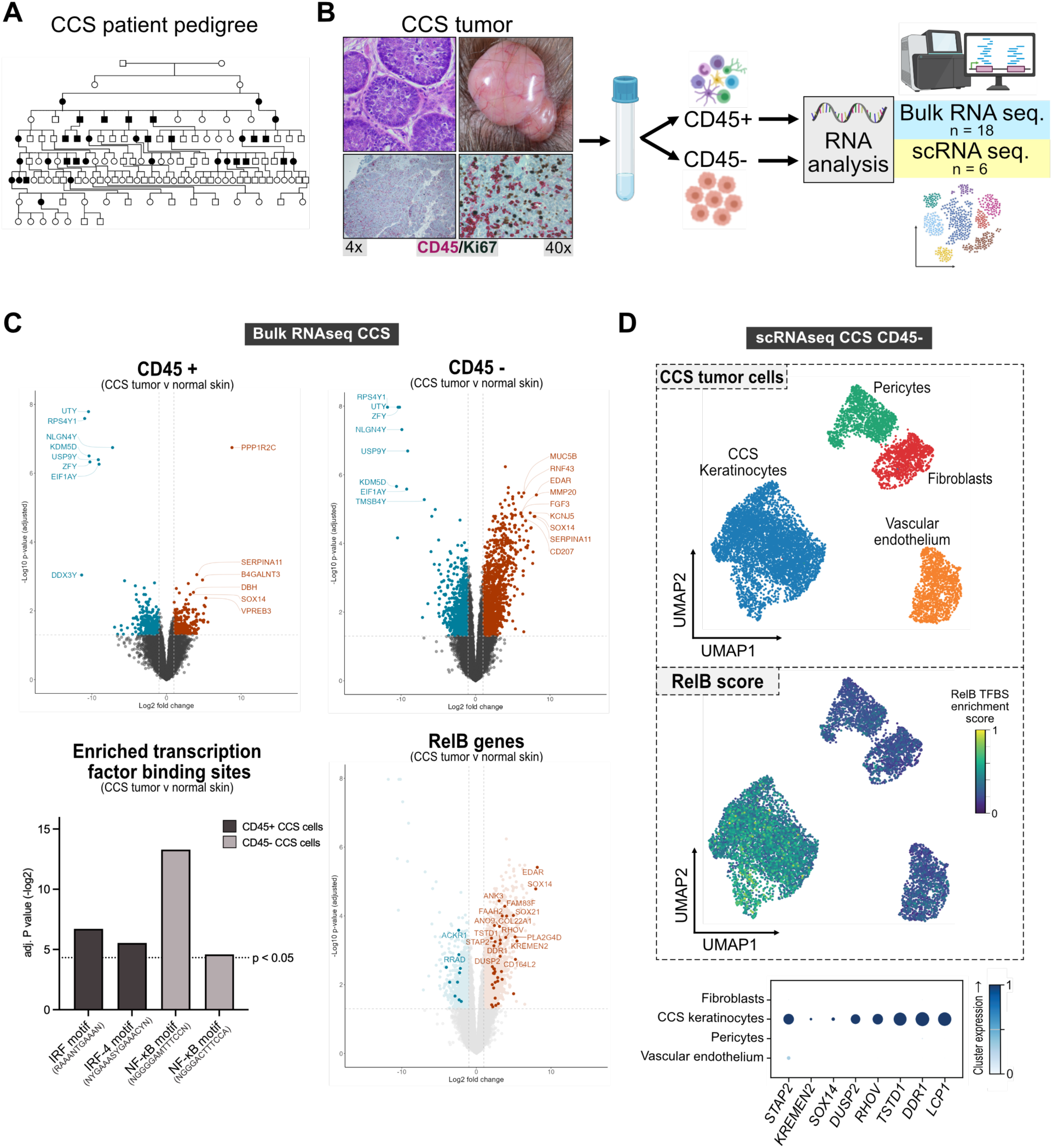
Expression of EDAR and EDA/EDAR target genes in CD45-enriched CCS transcriptomics. **A)** CCS patients develop tumors predominantly across the scalp and face from puberty onwards, with an autosomal dominant pattern of inheritance. **B)** Flow cytometry of enzyme disassociated CCS tumor and control skin suspensions was used to generate CD45^+/−^ single cell populations, that were used to generate bulk and single cell libraries for RNA sequencing. **C)** Differential gene expression between CCS tumors and normal skin was analysed in CD45^+/−^ fractions from bulk RNA sequencing data. The 500 genes with the highest increase in expression in CD45^+/−^ CCS tumor fractions were input to g:Profiler for functional enrichment analysis where statistically significant transcription factor motif matches were found (Benjamini-Hochberg corrected p-values). **D)** UMAP plot of unsupervised clustering of CD45^−^ cells from CCS tumours (n=3) according to similarity of transcriptome resulting in four clusters. Gene scoring per cell of the 49 DEGs with p52/RelB TFBS was performed and indicated as a RelB score overlaid onto a UMAP plot. Dot plot displaying mean gene expression for each cluster (colour) and percentage of cells expressing the marker within a cluster (dot size). UMAP Uniform Manifold Approximation and Projection; TFBS Transcription Factor Binding Site.

In sporadic cancers, *CYLD* is an important tumor suppressor, and is epigenetically repressed or mutated in cancers including leukemia (9), lymphoma (10), myeloma (11), neuroblastoma (12), and liver cancer (13). Recently, somatic truncating CYLD mutations have been frequently demonstrated in head and neck squamous cell carcinoma (14) and anal carcinoma (15), underscoring the relevance of studying the signalling consequences of *CYLD* loss of function mutations in cancer.

CYLD is a DUB with specificity for substrates tagged with Lys63- and Met1-linked polyubiquitin chains (16–18). In immune cells such as T cells for example(19), CYLD negatively regulates canonical NF-κB signalling (20, 21). CYLD deubiquitinates substrates that form part of signalling complexes linked to cell surface receptors including TNFR1, TCR, BCR and TLR2, as well as intermediate complexes such as IKK (22). Therefore loss of CYLD leads to RelA/p50 dimerization and transcription of target genes (20, 21, 23–26). Whole genome sequencing of CCS skin tumors has revealed biallelic inactivation of *CYLD* alone is an obligate event for tumor development, most frequently caused by copy neutral LOH of CCS skin tumors (27, 28). Bulk tumor transcriptomic studies to date of CCS skin tumors (cylindroma and related spiradenoma) have confirmed CYLD’s recognized roles in negatively regulating canonical NF-κB, Wnt/Beta-catenin (29, 30), tropomyosin receptor kinase (27), and TGF-beta signalling pathways (31). Additionally, sporadic spiradenoma and spiradenocarcinoma skin tumors that have a normal *CYLD* sequence are frequently driven by a somatic *ALPK1* hotspot mutation (p.V1092A). This mutation also enhances NF-κB activation, underscoring the importance of NF-κB signalling in the pathogenesis of CCS related tumors (32).

CYLD also negatively regulates non-canonical NF-κB signalling. Non-canonical NF-κB signalling, mediated via NIK and IKKα dimerization, leads to p100 to p52 processing, and p52/RelB mediated transcription of target genes (19). In murine keratinocytes, CYLD prevents the nuclear translocation of BCL3-p50/BCL3-p52 complexes via deubiquitination of BCL3 (33). Increased protein levels of RelB and p100 have been noted in B cells from both a Cyld knock-out (26) and a knock-in mouse model expressing a short splice variant of murine Cyld lacking exons 7 and 8 (34). A role for the non-canonical NF-κB signalling pathway has not however been previously demonstrated in CCS tumors keratinocytes (35, 36).

Dissection of the impact of truncating CYLD PVs on NF-κB signalling in CCS tumor cells have been hampered by the lack of mouse models that faithfully recapitulate the CCS skin tumor phenotype (33, 37). To overcome this, we used CCS patient tumors that are driven by natural mutagenic events affecting *CYLD* and offer a cellular signalling context relevant to the discovery of relevant therapeutic targets in CCS. Here, utilizing bulk and single cell transcriptomic and bulk proteomic profiling of CCS tumor fractions we demonstrate enrichment of non-canonical NF-κB signalling gene expression in CCS tumor keratinocytes. Ectodysplasin A receptor (EDAR) was the most highly expressed of the non-canonical NF-κB signalling gene signature, which in turn upregulates canonical NF-κB signalling in CCS tumor keratinocytes (38). To functionally investigate this, we developed a novel patient-derived primary CCS tumor keratinocyte enriched spheroid model. Targeting of non-canonical signalling in CCS spheroids by inhibiting IKKα using SU1644, a novel highly selective IKKα inhibitor, reduced EDAR expression and CCS spheroid viability at low micromolar concentrations. These data highlight IKKα as a candidate druggable target in CCS and support future studies of topical highly selective IKKα inhibitors in CCS patients.

## Results

### Non-canonical NF-κB signalling in CCS tumors drives EDAR overexpression

CCS skin tumors were first investigated to determine CD45 expression, and this demonstrated varying levels of CD45^+^ cell infiltration in the cylindroma and spiradenoma tumor microenvironment (**Figure 1B)**. As CD45^+^ immune cells include lineages which express NF-κB target genes (39), we depleted CCS tumor keratinocyte fractions of infiltrating CD45^+^ immune cells. To achieve this, we used CD45 labelling of enzymatically digested fresh CCS skin tumor cell suspensions followed by CD45^+^ depletion using FACS, an approach widely used to study CD45^−^ tumor cells in cancer (40) (**Figure 1B)**. Bulk RNAseq analysis was performed on flow sorted CD45^+^ (n=9) and CD45^−^ (n=9) fractions, with additional samples sequenced for quality control purposes (whole CCS tumor (n=6) and control skin (n=3) snap frozen tissue, CD45 labelled unsorted CCS single cell suspensions (n=5) and normal skin controls (n=4); see **Supplementary Table 1** for full list of patient samples and principal component analysis (**Supplementary Figure 1**).

CCS tumor CD45^−^ cells expressed a cytokeratin profile recognized in CCS tumors (6), including increased expression of *KRT8*, *KRT17* and hair follicle cytokeratin *KRT74* (adjusted *P*-value < 0.05) (**Supplementary Figure 2**A), confirming CCS CD45^−^ fractions were enriched for CCS tumor keratinocytes. Functional enrichment analysis of differentially expressed genes in CCS CD45^+^ and CD45^−^ cell fractions compared to respective control skin fractions was performed using g:Profiler (42). To identify up-regulated genes in CCS fractions with NF-κB transcription factor binding sites (TFBS), the TRANSFAC PWM library and MATCH (43) tool was used. The most significant match was in the CD45^−^ cell fraction, where the p52/RelB heterodimer motif (NGGGGAMTTTCCNN) was significantly enriched (adjusted p<0.0001), indicative of upstream non-canonical NF-κB signalling (**Figure 1C**). 798 putative genes with at least one p52/RelB TFBS match (motif GGGGNTTTCC) +/−1kb from the transcriptional start site (TSS) were identified from the TRANSFAC PWM library. Next, we filtered for genes with both a p52/RelB TFBS, and which were differentially expressed in CD45^−^ CCS tumor cells (FC >2, adjusted *P*-value < 0.05 (Benjimini-Hochberg multiple test correction). Out of 1767 DEGs, 49 had a p52/RelB motif +/−1kb from the TSS, of which *EDAR* was the most differentially expressed (8.2x Log2FC in tumor compared to control; **Supplementary Figure 3, Supplementary Table 2**). EDAR is recognized to drive canonical NF-κB signalling itself in hair follicle development and is overexpressed in Wnt/beta-catenin mediated breast cancers (44, (45)), which led us to assess genes it modulates in CCS tumors. We analyzed differential expression of a literature curated set of EDAR signalling pathway genes (46–49) in the bulk CD45^−^ cell data. Consistent with active EDAR signalling, sixteen candidate EDAR regulated genes were differentially expressed in CCS tumor cells compared to control, which included: Kringle containing transmembrane protein 2 (*KREMEN2*), Wnt family member 10B (*WNT10B*) and TNF superfamily receptor family 18 (*TNFRSF18*, syn. *GITR*) (**Supplementary Figure 2B**).

To confirm the p52/RelB TFBS genes differentially expressed in the CD45^−^ CCS tumor cells originated from CCS tumor keratinocytes, we generated a “RelB score” from these genes in the bulk data that we explored at single cell resolution. First, we characterized the cell populations present in the CD45^−^ CCS samples using single cell RNAseq of CCS CD45 labelled tumors cells (n=3 samples; K=7349 cells). In the CD45^−^ fraction, we detected CCS tumor keratinocytes (*KRT14, KRT17*), fibroblasts (*LUM, CFD),* pericytes (*RGS5, GJA4*) and vascular endothelial *cells* (*PECAM1, ESAM*) (**Figure 1D).** CCS tumor keratinocytes were found to have an increased “RelB score” compared to other clusters (**Figure 1D)**, and increased expression of p52/RelB target genes identified in bulk transcriptomic data sets including *STAP2, KREMEN2, SOX14, DUSP2, RHOV, TSTD1, DDR1, and LCP1*.

### CD45^−^CD200^+^ CCS tumor cells express IKKα, an upstream regulator of non-canonical NF-κB signalling

To validate our observation of deregulated non-canonical and canonical NF-κB signalling in CCS tumor keratinocytes we characterized the CCS tumor proteome with DIA (data independent acquisition) LC-MS (liquid chromatography-mass spectrometry). Given our patient guided therapeutic goal of developing a medical treatment that could target early and pre-clinically apparent tumors in genotyped CCS patients, we sought to enrich for CCS tumor initiating cells (TICs). We used CD200, a highly expressed TIC marker (50) in CCS tumor cells (3), to enrich the CCS CD45^−^ cell fraction in CCS tumors (**Figure 2A**). We compared additional flow sorted CD45^−^CD200^+^ CCS tumors (n=5) and CD45^−^ enzymatically-split epidermis alone from healthy skin (n=5). The number of protein identifications (IDs) in the tumor proteome (mean = 6666) compared to healthy epidermis (mean = 1891) revealed a remarkable increase in the proteome complexity of CCS tumors withstanding median normalization (**Supplementary Figure 4**), and showed differential clustering by PCA (**Figure 2B**). We detected a total of 7204 proteins across all samples, with 2677 tumor specific proteins, and 3245 proteins detected in at least three replicates in both conditions. CD45^−^CD200^+^ cells demonstrated an increase in hair follicle specific keratins (**Figure 2C**). We explored the expression levels of NF-κB pathway proteins and proteins encoded by NF-κB target genes in CD45^−^CD200^+^ cells. We detected increased protein expression of regulators and components of the non-canonical NF-κB pathway including TRAF2 and TRAF6, CHUK/IKKα, p52/NFKB2 and RELB in CCS cells compared to control skin. Consistent with EDAR having a p52/RELB motif, increased expression of EDAR was the only cell surface receptor known to regulate NF-κB detected in CCS tumors (**Figure 2C-D**). Evidence of canonical EDAR/NF-κB signalling pathway regulators and components was also evident with increased expression of MAP3K7/TAK1, IKBKG, IKBKB, p65/RELA and proteins encoded by NF-κB target genes including BCL2, XIAP and RIPK1. In these CD45^−^CD200^+^ cells, there was an increase in beta-catenin (CTNNB1 3.14x Log_2_FC (**Supplementary Figure 5**) and a reduction in glycogen synthase kinase-3 beta (GSK3B −3.2x Log_2_FC), consistent with beta-catenin stabilization, and prior work showing nuclear beta catenin in CCS cells (30).The detection of EDAR overexpression in CCS in the context of active Wnt/beta-catenin signalling is consistent with the synergy reported in luminal breast cancer (45).

**Figure 2.**
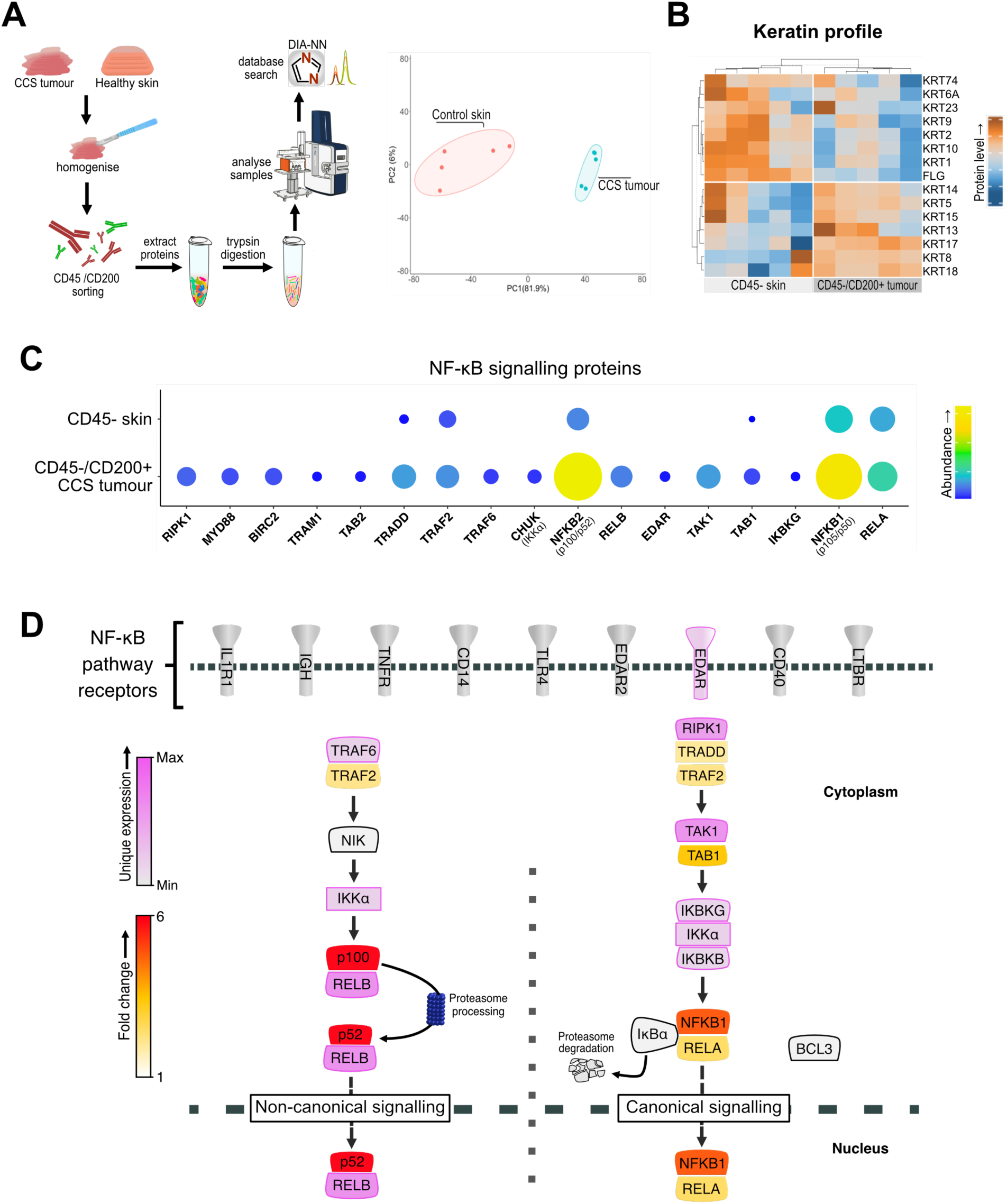
Proteomic analysis of CD45^−^CD200^+^ CCS tumour cells and CD45-normal skin reveal increased EDAR and NF-κB signalling pathway members. **A)** CCS tumour and enzyme split epidermis from healthy skin samples were processed to enrich CD45^−^/CD200^+^ tumour cells (CCS tumour keratinocytes) and CD45^−^ skin cells (healthy keratinocytes) respectively for proteomic analysis. Principal component analysis revealed distinct proteomes. **B**) Keratin profiling of CCS tumour keratinocytes and healthy keratinocytes by hierarchical clustering of highly expressed keratin genes and filaggrin. **C**) The abundance (size and colour) of NF-κB signalling proteins in CCS tumour keratinocytes compared to CD45^−^ skin. **D**) Detected NF-κB signalling pathway proteins are projected onto a signalling diagram, showing the proteins uniquely detected in CCS tumours (pink) and the fold change increase (yellow to red). Grey shaded proteins were not detected.

### CCS tumors and tumor spheroids demonstrate dysregulated NF-κB signalling with increased p100/p52 and RelB expression

To functionally investigate our findings given the lack of a mouse model, we developed a novel CCS spheroid culture model from primary tumor cells derived from fresh surgical samples of CCS skin tumors (**Figure 3A Supplementary Figure 6, Supplementary Table 1**). CCS skin tumors were enzymatically dissociated and cultured in defined keratinocyte serum free medium (0.05 mM Ca^2+^) for 24 hours, before spheroids were formed in Aggrewell plates (StemCell UK) at increased calcium concentrations (1.2mM Ca^2+^), known to facilitate primary keratinocyte spheroid formation without inducing differentiation (51). CCS keratinocyte enrichment was confirmed by expression of truncated CYLD (trCYLD; c.2460delC) and lack of expression of full length CYLD (flCYLD), which was used to select samples to investigate NF-κB keratinocyte signal transduction as indicated below. CCS spheroids recapitulated the same pattern of cytokeratin expression to that seen in *in vivo* (**Figure 3A**). In addition, CCS spheroids proliferated in culture, as indicated by expression of Ki-67, and demonstrated evidence of previously reported overexpressed *in vivo* proteins including beta-catenin (30).

**Figure 3.**
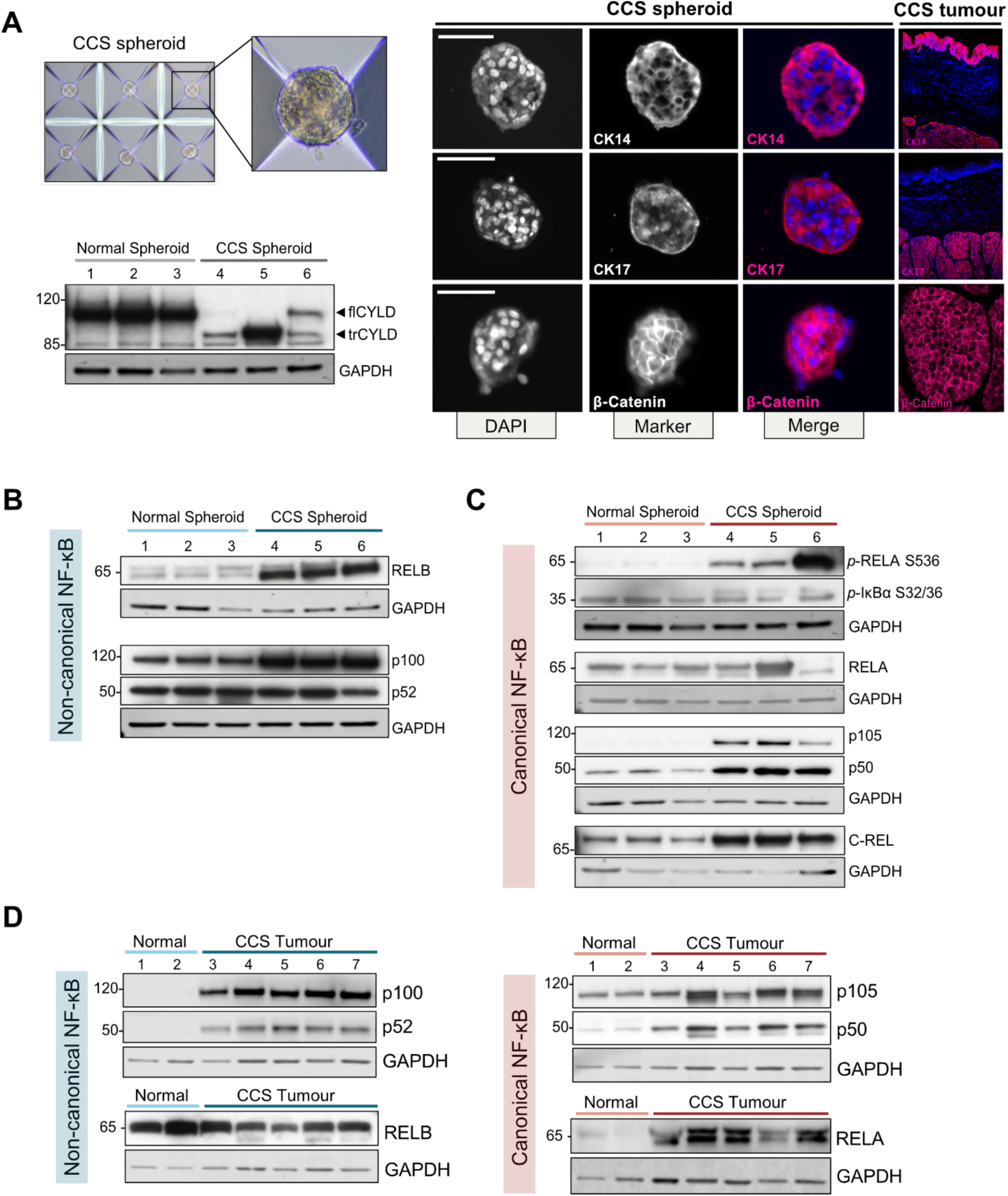
CCS tumours and tumour spheroids demonstrate dysregulated NF-κB signalling with increased p100/p52 and RelB expression. **A)** CCS tumour spheroids are an *in vitro* model for CCS tumours which recapitulate expression of CCS tumour proteins and express truncated CYLD (trCYLD). Images are representative of ≥ 5 spheroid generation experiments, 20x magnification, scale bar = 50 mm. **B)** CCS tumour spheroids demonstrate increased expression of non-canonical and **C)** canonical NF-κB members. **D)** CCS whole tumour lysates display dysregulated NF-κB signalling compared to normal skin controls. Lane numbers 1-6 in (A),(B) and (C) indicate respective protein lysates used for these panels.

Both canonical and non-canonical NF-κB pathways were dysregulated in CCS spheroids relative to normal primary keratinocyte spheroids (**Figure 3B-C**). Unlike normal whole skin (**Figure 3D**), p100 to p52 processing was already at a high level in the normal spheroids, possibly due to the increased proliferative state in culture compared to normal skin. In CCS spheroids there was an increase in p100 and RelB protein, and p52 levels were comparable to the control primary keratinocyte spheroids. In the canonical pathway c-Rel and p105/p50 protein were increased in CCS spheroids. Moreover, increased levels of RelA phosphorylated at serine 536 suggest the canonical signalling pathway is also active in tumor keratinocytes (52). We also investigated a lysate sample from CCS spheroids (**Figure 3B,C -** Lane 6) where tumor microenvironment (TME) cells had persisted, as evidenced by presence of flCYLD, and found that canonical NF-κB signalling (evidenced by increase in RelA phosphorylation) was altered compared to CCS spheroids without TME cells where this was lower (**Figure 3B,C -** Lanes 4,5).

Finally, to compare the spheroids to CCS patient tumours, we studied these same proteins in snap frozen whole CCS tumour protein lysates and normal control skin (**Figure 3D**). Here, we also found increased p100 and p52 protein levels and similar RelB levels in CCS tumor lysates, compared to normal skin. Increased RelA and p50 protein levels were also present. These data support CCS spheroids as a faithful *in vitro* model of CCS, and allowed us to proceed to study the effect of targeting non-canonical NF-κB signalling by inhibiting IKKα.

### Selective IKKα inhibition reduces CCS tumor spheroid viability and is associated with a reduction in truncated CYLD and non-canonical p100 to p52 processing

NIK and IKKα phosphorylate p100 leading to its ubiquitination and processing to p52 by the proteasome, and in addition p100 inhibits RelB nuclear localization (53). IKKα is upregulated in CD45^−^CD200^+^ CCS tumor cells **(Figure 2C**), and is not itself shown to be mutated in CCS from previous tumor sequencing studies (28). Therefore, selectively targeting IKKα has the potential to reduce non-canonical NF-κB signalling and be a novel therapeutic strategy for CCS. However to date, highly IKKα selective inhibitors have been lacking. As part of an ongoing drug discovery program to develop selective IKKα inhibitors that demonstrate low affinity for IKKβ, we have recently developed SU1644. SU1644 demonstrated selective IKKα target engagement in a metastatic prostate cancer cell line (PC3m) using pharmacodynamic markers (p100 phosphorylation; RelB nuclear translocation) [IC_50_ = 0.05 µM], with no activity against IKKβ markers evident at 10 µM (data not shown). SU1644 also had an excellent kinase selectivity profile when assessed against a representative kinome sample (100 kinases: **Supplementary Figure 7**). The development of this selective IKKα molecule and a related series has recently been patented (PCT/GB2023/051242). Topical delivery directly to the skin of SU1644 itself is attractive because of its high hepatocyte clearance, (220 μL/min/10^6^ cells; t_1/2_ = 6 min), which would lead to low systemic exposure and thus avoid perturbation of non-canonical NF-κB signalling in other tissues (data not shown). We used the highly isoform-selective IKKα inhibitor, SU1644, and found that this had an IC_50_ of 2.74μM in our CCS spheroid model (**Figure 4A**). We targeted IKKβ specifically with TPCA1 (**Figure 4B**) and both IKKα and IKKβ with BMS-345541 (IC_50_ values are 0.3 and 4.0 μM for IKKβ and IKKα respectively) (**Figure 4C**) which both showed higher IC_50_ values of 5.76 μM and 5.56 μM, respectively, in our CCS spheroid model. We studied the effect of selective IKKα inhibition using SU1644 on truncated CYLD (trCYLD) in CCS. SU1644 reduced trCYLD levels at 1-3μM concentrations in CPCC cultures from patients with a truncating PV and a splice site PV (**Figure 4D, Supplementary Figure 8**). SU1644 also reduces p100 processing and p52 levels consistent with a decrease in non-canonical NF-κB signalling (**Figure 4E**). CCS spheroid *EDAR* expression was also significantly reduced after SU1644 treatment, confirming its regulation by the non-canonical NF-κB pathway implied above (**Figure 4F**). Together, these findings identify IKKα inhibition as a novel candidate for CCS tumor prevention.

**Figure 4.**
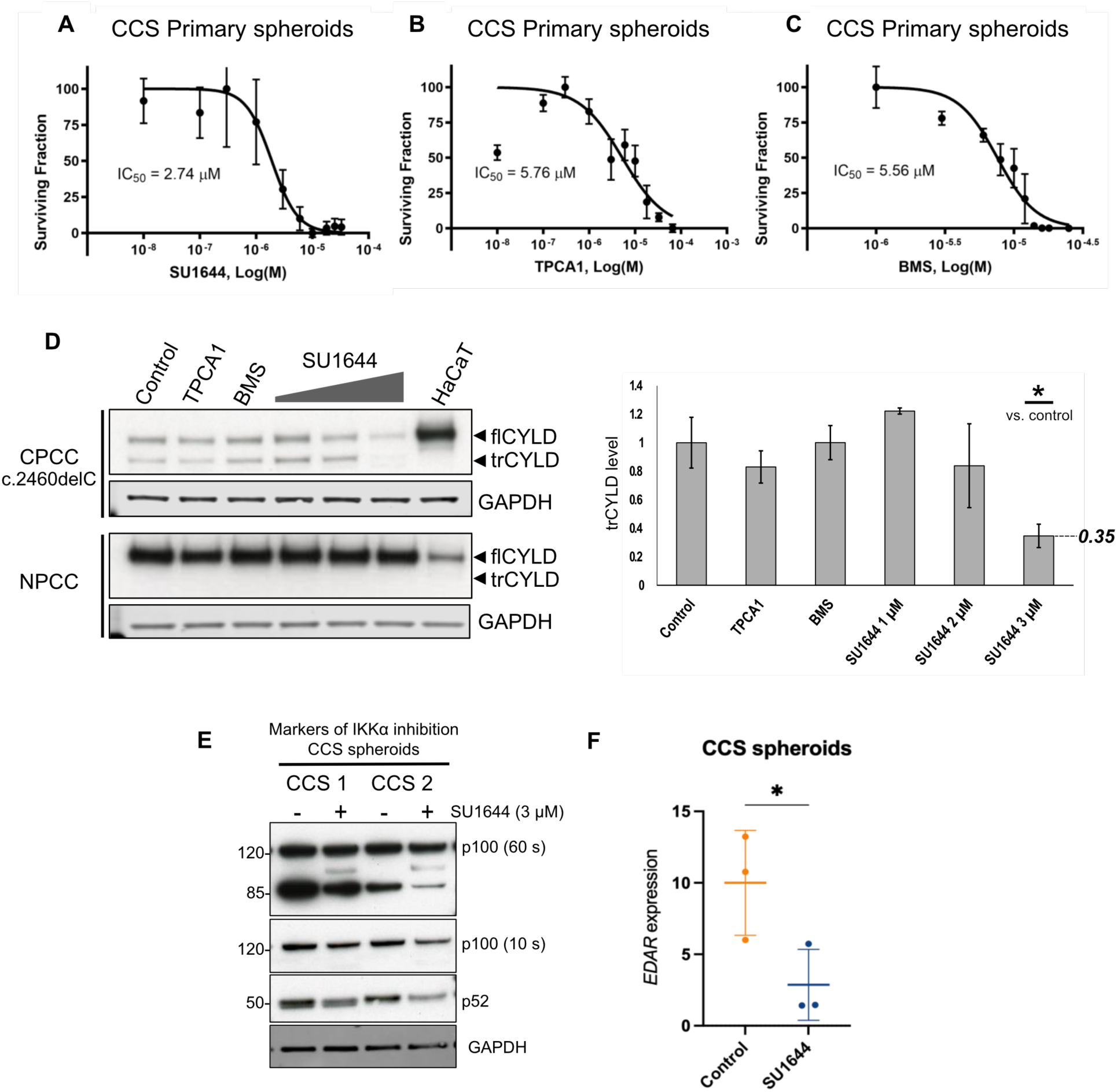
Targeting IKK⍺ with the highly selective IKK⍺ inhibitor SU1644 results in reduced CCS spheroid viability, and reduced trCYLD and EDAR expression. CCS spheroid dose-response to small molecule IKK inhibitors assessed by CellTiter-Glo luminescent viability assay after treatment with **A**) SU1644 targeting IKK⍺ **B**) TPCA-1 targeting IKKβ or **C**) BMS-345541 targeting IKK⍺ and IKKβ. **D**) Levels of trCYLD (truncated CYLD) and flCYLD (full length CYLD) in CCS primary cell culture (CPCC) and normal primary cell culture (NPCC) detected by immunoblotting after 24 h treatment with IKK inhibitors. HaCaT lysate is included as a positive control for flCYLD molecular weight only. TPCA1 and BMS; BMS345541 were used at concentrations of 6 μM. **E**) p100 to p52 is reduced in CCS tumour spheroids after 24 h SU1644 treatment. **F**) EDAR RNA expression measured by real time quantitative polymerase chain reaction (RT-qPCR) is reduced in CCS spheroids following 24 h 3 μM SU1644 treatment. Significance is indicated as * = p < 0.05 after Students t-test.

## Discussion

CCS is a progressive condition which lacks medical therapies, and mechanistic understanding of oncogenic drivers in tumor keratinocytes are a prerequisite for the development of novel treatments. By studying patient samples with naturally occurring *CYLD* PVs, this work using complementary transcriptomic and proteomic methods, offers insight into non-canonical and canonical NF-κB signalling in CCS tumors. Increased p100 and p52 levels *in vivo* and in CCS spheroids are a hallmark of non-canonical signalling. Since the p100 precursor protein also inhibits RelB nuclear localization, NIK and IKKα which mediate p100 processing are attractive druggable targets (53). Here, we demonstrate that the targeting of IKKα with SU1644 in the non-canonical NF-κB pathway reduces *EDAR* expression and cell viability in patient derived CCS spheroids.

EDAR overexpression has recently been found to synergize with increased beta-catenin signalling in mouse mammary cancer models towards driving tumorigenesis (45). Precise spatiotemporal expression of EDAR/NF-κB signalling in human development is crucial for hair follicle, skin appendage and tooth development (54). Loss of function mutations in these genes lead to reduction in EDAR/NF-κB signalling and ectodermal dysplasia phenotypes with absent hair, sweat glands and teeth (55, 56). We propose that together with activated Wnt/Beta-catenin signalling (29, 30), EDAR signalling in CCS promotes CCS tumor keratinocyte viability. Expression of antiapoptotic targets of canonical NF-κB signalling such as BCL2, previously described in these tumors (27) suggest that the progressive growth that results in the exceptionally large CCS tumors (1) may be mediated by resistance to apoptosis.

Delineation of the upstream events that lead to increased p100/p52 processing remains to be determined, but we provide data that implicates a hypomorphic trCYLD protein. Murine studies have established that the phenotype associated with CYLD truncation or expression of alternate short isoforms is distinct from that seen in complete CYLD ablation (22). In humans, CCS tumors are predominantly driven by truncating mutations, suggesting the hypomorphic protein in the absence of full length CYLD is likely to be a key driver. Truncated CYLD expression was seen to be reduced following IKKα inhibition, and this may relate to IKK’s recent role in the regulation of mRNA stability (57, 58). This was associated with reduced spheroid viability and p100 to p52 processing. The reduction of truncated CYLD may also reflect the selective killing of CCS tumor keratinocytes by SU1644, and the IC_50_ for tumor cells may be lower than reflected in **Figure 4A**. Taken together, these suggest that trCYLD interferes with regulation of non-canonical NF-κB signalling. Given the multiple targets of CYLD, this is likely to be mediated by more than one substrate. It remains to be understood how the expressed CYLD protein in infrequent (2) missense *CYLD* PV carriers abrogates signalling, however work to understand the crystal structure of predicted truncated variants (59) suggest that analysis of tertiary protein structures may reveal how conformational changes from missense PVs may also impact on interference with regulation of NF-κB signalling.

We studied CD45^−^CD200^+^ CCS tumor cells as CD200 is a marker of hair follicle stem cells, and are proposed tumor initiating cells, as seen in other skin tumors such as basal cell carcinoma (38). Using an unbiased proteomic approach, we were able to determine the presence of increased non-canonical and canonical NF-κB signalling in CD200^+^CD45^−^ CCS cells. This is especially important in the development of a “pre-emptive” topical treatment applied to at-risk skin for genotyped CCS patients, as IKKα inhibition can potentially kill off early, pre-clinically apparent cells. CCS is particularly suitable for topical delivery of inhibitors of IKKα, circumventing potential side effects such as those seen with systemic delivery of IKKbeta inhibitors (60). These data suggest the selective targeting of IKKα may lead to translational advances for patients living with CCS.

## Materials and Methods

### Statement regarding sex as a biological variable

Our study examined male and female samples, and similar findings are reported for both sexes.

### Study Approval

Research ethics committee approval was obtained from the Hartlepool Research Ethics Committee and North East – Newcastle & North Tyneside 1 Research Ethics Committee for this work (REC Ref: 06/Q1001/59; 08/H0906/95+5; 19/NE/0004). Skin samples were obtained from patients with signed, informed, consent and details of samples are shown in **Supplementary Table 1**. Written informed consent was received for the use of the photographs and that the record of informed consent has been retained.

## Spheroid primary cell culture

Aggrewell400™ 6-well plates containing 7000 microwells per well (Stemcell Technologies) were prepared with 2 ml/well anti-adherence rinsing solution (Stemcell Technologies 07010), centrifuged at 1300 x g for 5 min and rinsed with 37°C cell culture medium (complete KSFM for CCS primary cells or complete Epilife for normal primary cells). Cell culture medium supplemented with 1.5 mM CaCl_2_ and 10 μM rho kinase inhibitor (ROCKi) was added at 2 ml/well.

Two confluent 75 cm^2^ flasks of 2D primary cells were trypsinised, pooled, and resuspended in 6 ml media/1.5 mM CaCl_2_/10 μM ROCKi. One ml of cell suspension (containing 5 × 10^5^ - 2 × 10^6^ cells) was added to each well of the 6 well Aggrewell and gently mixed. After centrifuging the Aggrewell plate at 100 x g for 3 min, wells were topped up to 5 ml with media containing 1.5 mM CaCl_2_ and 10 μM ROCKi. Media was changed after 48 h by aspirating and replacing with 2.5 ml media containing 1.5 mM CaCl_2_.

### RNAseq library preparation

RNA was extracted from FACS sorted samples with the RNeasy Micro Kit (Qiagen 74004) and bulk tissue with the Allprep DNA/RNA/miRNA Universal Kit (Qiagen 80224) following the manufacturers’ protocols. TapeStation (Agilent) RNA quality control and stranded preparation with the NEB Nextera Low Input RNA Library Prep Kit was performed. Libraries were single-end sequenced using an Illumina NextSeq 550 system. scRNA sequencing library generation was performed utilizing a 10X Genomics Chromium Single Cell *5*’ GEM Library generation kit according to manufacturer protocols. Libraries were pair-end sequenced using a S1 flow cell on an Illumina Novaseq system.

### Bioinformatic analysis

Bulk RNA sequencing data quality was assessed with FastQC and reads were aligned using the splice-aware aligner program STAR (61). The aligned sequencing reads for each sample were mapped to genomic features and counted using the Python package HTSeq (62). Count data was filtered to remove genes with a total read count <15, normalized by trimmed mean of M values (TMM) method in the Bioconductor package edgeR (63) and then transformed by voom (64) using the limma R package (65). Differential gene expression analysis was carried out using the package DeSeq2 (66). Graphical analyses were generated using ggplot2 (67). Functional enrichment analysis of differentially expressed genes was performed with the webserver g:Profiler (42). Single cell RNA sequencing BCLs were processed using Cellranger 3.0.1 and aligned to Cellranger reference GRCh38 3.0.0. Cells with fewer than 200 genes or >10% mitochondrial transcripts were filtered. Batch correction by sequencing lane was performed using Harmony (harmonypy version 0.0.9). Gene scoring per cell of the 49 DEGs with p52/RelB TFBS (Fig 1D) was performed using the tl.score.genes module in Scanpy. This calculates average expression of the provided gene list subtracted by a random sampling of background genes.

### Proteomics sample preparation and analysis

FACS sorted samples were prepared for proteomics analysis by lysis in 5% SDS, 8 M urea, 100 mM glycine pH 7.55 followed by concurrent reduction with 20 mM tris (2-carboxyethyl) phosphine hydrochloride (TCEP) and alkylation with 20 mM Iodoacetamide for 30 min at 37 °C in the dark. Samples were acidified with phosphoric acid to 1% v/v. Proteins were precipitated with 90% Methanol, 10% TEAB v/v, and the resulting mixture added to ProtiFi micro S-Trap column following the manufacturer’s protocol. Briefly, samples were passed through a column by centrifugation at 4,000 x g for 1 min and washed with 90% Methanol, 10% TEAB v/v 5 times by centrifugation. Column bound proteins were digested with 1:20 trypsin at 47 °C for 2 h. Resulting peptides were serially eluted with 40 μL of 50 mM TEAB pH 8.0, 0.2% FA in water and 0.2% FA in 50% acetonitrile, 50% water. Eluates were dried by vacuum concentration and resuspended in 0.1% formic acid in water.

500 ng of digest was analyzed on a Bruker TIMS-TOF HT mass spectrometer in line with the EvoSep One LC system, as previously reported (68). Samples were separated by elution in a gradient of MS solvent A (0.1 % formic acid in HPLC water) and MS solvent B (0.1 % formic acid in HPLC grade acetonitrile), by the manufacturer-defined Evosep One Whisper 20 samples per day method. The mass spectrometer was operated in positive-ion, data independent acquisition parallel accumulation serial fragmentation (DIA-PASEF) mode. Precursor ion scans (full scan, MS1) were performed in the range of 100-1,700 *m/z*, for DIA-PASEF (MS2 scans), mass and IM ranges were from 300-1200 *m/z* and 0.6-1.4 1/*K_0_*, DIA-PASEF was performed using 16 variable width IM-*m/z* windows, designed using py_diAID (69), details are previously published (68). Collision energy was applied as a linear gradient from 20-59 eV to precursors with mobilities from 0.60-1.60 V.s/cm2. The ramp and accumulation time were set to 100 ms, with a ramp rate of 9.42 Hz, the total cycle time was ∼1.8 sec.

Data files were searched against an in-silico digest of SwissProt *Homo sapiens* database performed in DIA-NN 1.8.1(70). Analysis of protein groups intensities was performed in the R statistical computing environment v4.3.0; including data filtering, clustering, principal component analysis, t-tests and graphics generation. The packages employed included limma (65), tidyverse (71) and ggplot2 (67).

### Statistical analysis

Statistical analysis was performed in R computing environment, utilizing base R functions and previously mentioned packages to perform statistical tests and produce visualizations. RT-qPCR statistical analysis was performed in Graphpad Prism software. Graph error bars indicate standard deviation. Statistical tests used and *P* values are noted in figure legends.

Further materials and methods are detailed in a supplementary file.

## Supporting information

Unedited Western blot images are available in a separate file

## Acknowledgments

We are indebted to the patients and families who took part in this study.

## Author contributions

Conceptualization: NR, NDP

Preliminary experiments: ES, EA, NN

Clinical sample acquisition: NR, KH, JI

Investigation: KH, JI, NS, AP, GC, CL, CJL and AMF

SU1644 compound design and development: AP, GB, CL, SPM

Bioinformatic analysis: KH, JI, SC, GR

Visualization: JI, KH

Supervision: NDP, II, SPM, NR

Writing—original draft: KH, JI, NR, NDP, SPM, MS, DA

Writing—review & editing: All authors

## Data availability

Bulk RNA sequencing data is available at EGA - EGAD50000000645 and scRNA sequencing is available at EGAD50000000702. The files are directly available here for reviewer access https://www.dropbox.com/scl/fo/pkeo6ve37owgki754tumm/AL1axHeZ_Tz0XT15dEOqOjk?rlkey=cx2yjdjyd4xvky06spwlggnga&st=pl3ja1sc&dl=0 The mass spectrometry proteomics data have been deposited to the ProteomeXchange Consortium via the PRIDE [1] partner repository with the dataset identifier PXD053466. The proteomics data is uploaded to ProteomeXchange with the following details for the reviewer to access: Username, Password: 9jm9Llgui4Cc. All data are available in the main text or the supplementary materials.

Unedited blot images are available in a separate file.

