## Supplementary Figures, Tables and Materials and Methods for "Targeting non-canonical NF-κB signalling in CYLD cutaneous syndrome by selective inhibition of IκB kinase alpha"

### Supplementary Materials and Methods

#### Cell culture

CCS tumors were collected in Dulbecco's modified eagle medium (DMEM) with 10% fetal bovine serum (FBS) immediately after surgery, transported to a class II cell culture hood, cut into ~1mm$^3$ pieces and transferred into 15 ml centrifuge tubes. One 75 cm$^2$ flask per tumor was coated with collagen (R011K Gibco) and incubated at room temperature during sample preparation. After centrifugation at 900 x g and decanting the supernatant, tumors were digested in 5 ml 2.5% trypsin for 40 min at 37°C with gentle agitation every 20 min. Tumors were centrifuged at 900 x g to allow aspiration of 3 ml trypsin. The pellets were incubated in 5 ml collagenase (1 mg/ml) for 70 min at 37°C with gentle agitation every 20 min. The dissociated tumor suspensions were transferred to 50 ml centrifuge tubes and 3 volumes of DMEM added before passing through a 40 um filter into a new 50 ml tube. Each quarter of the cell suspension was transferred to a 15 ml centrifuge tube and centrifuged at 900 x g. The resulting pellet was resuspended in complete defined keratinocyte serum free medium (DKSFM Gibco 10744019)/0.2% penicillin-streptomycin (5000 units/ml)/1% L-glutamine (200 mM)/0.04% Amphotericin B (250 µg/ml) and seeded in a collagen coated 75 cm$^2$ flask for continued culturing. Alternatively, the pellet was resuspended 10% dimethylsulfoxide (DMSO)/FBS and cryopreserved at -80°C. Cell cultures were maintained in complete DKSFM supplemented with 10 µM rho kinase inhibitor (Y-27632 Tocris) and passaged when ~80% confluent.

#### Normal skin primary cell culture

Immediately after surgery for non-CCS related conditions, donated skin was collected in complete DMEM with Dispase II (1 unit/ml) and incubated on a roller at 4°C overnight. One 25 cm$^2$ flask per sample was coated with collagen (R011K Gibco) and incubated at room temperature during sample preparation. The epidermis was peeled from the dermis with fine forceps, transferred to a 15 ml centrifuge tube containing 5 ml 0.05% Trypsin-EDTA and incubated for 30 min at 37°C with gentle agitation by pipetting every 15 minutes. After adding 5 ml complete DMEM and centrifugation at 300 x g for 5 min, the supernatant was aspirated and the pellet resuspended in complete Epilife medium (Gibco MEPI500CA)/0.2% penicillin-streptomycin (5000 U/ml)/1% L-glutamine (200 mM)/0.04% Amphotericin B (250 µg/ml). Cells were seeded onto collagen coated flasks and passaged when 80% confluent. Cells were maintained in complete Epilife medium supplemented with 10 µM Rho kinase inhibitor (Y-27632 Tocris).

#### 2D Cell passage and harvesting for protein

Cells were passaged by washing with phosphate-buffered saline (PBS), incubating in versene (Gibco 15040-066) for 5 min at 37°C followed by 0.05% Trypsin-EDTA at 37°C until dissociated. After quenching with complete DMEM, cells were centrifuged at 900 x g and resuspended in 1 ml PBS. Flasks were reseeded with 20% of the cell suspension and the remaining cells were transferred to a microcentrifuge and centrifuged at 17,000 x g for protein extraction. The supernatant was completely aspirated and pellets were frozen on dry ice immediately and stored at -80°C.

**Harvesting spheroids for protein**

On day 4 of spheroid culture, ¾ media was removed from each well and the remaining media pipetted firmly to dislodge the spheroids from the microwells. The spheroid containing media was passed through a 40 um filter into a 50 ml tube to remove single cells. Cell culture medium (1 ml) supplemented with 1.5 mM $CaCl_2$ and 5 mM EDTA was used to wash each well twice and each wash was passed through the filter. The filter was then inverted and spheroids were recovered by rinsing with 5 ml media/1.5 mM $CaCl_2$/5 mM EDTA into a new tube. The spheroid suspension was centrifuged at 2000 x g for 5 min and the media aspirated. The resulting pellet was frozen immediately on dry ice and stored at -80°C for downstream protein extraction.

**Cell viability assay**

Primary cells were seeded in 96 well spheroid microplates (Corning 4520) at 30,000 per well in 280 µl complete media supplemented with 1.5 mM CaCl2. After 3-7 days, spheroids were drugged at the range of concentrations indicated. After 72 hours incubation with TPCA1/BMS345541 or 48 hours incubation with SU1644, cell viability was assessed with the CellTiter-Glo® Luminescent Cell Viability Assay (Promega). Luminescence was recorded with the Varioskan™ LUX microplate reader. Dose response curves were generated using GraphPad PRISM version 9. Error bars are standard deviation of the mean of three replicates wells for each concentration. Three biological repeats were performed for each experiment, with representative data and mean $IC_{50}$ values shown in the figures.

**Protein extraction**

Tissue curls were cut at 30 µm on a cryostat from snap frozen CCS tumour or normal skin. Tissue was lysed in 150 µl Novagen's PhosphoSafe™ Extraction Reagent (EMD Millipore) or 8M Urea lysis buffer (8M Urea, 50 mM Tris pH 8, 300 mM NaCl, 50 mM Na2HPO4, 0.5% NP-40) and homogenised in Precellys® CK Mix 2 ml bead tubes. Samples were centrifuged in the bead tubes at 4°C, 15,000 x g for 5 min to reduce bubbles then transferred to microcentrifuge tubes. After centrifuging again at 4°C for 15 min at 15,000 x g, samples were sonicated 3 times for 15 s at 30% amplitude. Samples were centrifuged at 4°C for 20 min at 15,000 x g and the supernatant was transferred to a new microcentrifuge tube. Protein was extracted from spheroid and 2D cell pellets by sonication for 3 x 15 s at 30% amplitude in 40–100 µl 8M Urea lysis buffer. Protein concentration was measured with the Pierce™ BCA protein assay kit.

**Immunoblotting**

Samples were prepared with NuPAGE™ lithium dodecyl sulphate (LDS) loading buffer and dithioreitol (DTT) sample reducing agent (Invitrogen) to 0.5 µg/µl protein. Samples were heated at 70°C for 10 min, loaded on NuPAGE Tris-Acetate or Bis-Tris precast gels in the appropriate running buffer (NuPAGE Tris-Acetate, MES or MOPS sodium dodecyl sulphate (SDS) running buffer, Invitrogen) and separated with the NuPAGE gel electrophoresis system before transfer onto PVDF membranes with the Bio-Rad Transblot Turbo semi-dry transfer system. PVDF membranes were blocked in Tris buffered saline with Tween-20 detergent (TBST) (20 mM Tris pH 7.6, 120 mM NaCl, 0.1% Tween 20) and 5% w/v fat-free milk or 3% w/v BSA for 1 h. Membranes were subsequently incubated in primary antibody solution overnight at 4°C and secondary antibody conjugated to horseradish peroxidase (HRP) (Cell signalling) or a fluorescent dye for 1 h at room temperature (see Table 2-8 for antibodies). Washes between incubations were performed in TBST for 3 x 10 min. HRP enzyme activity was detected with enhanced chemiluminescent (ECL) substrate. Fluorescence was detected with the LICOR Odyssey® CLx imaging system.

**Antibodies used for immunoblotting.** CST, Cell Signaling Technology; Thermo, ThermoFisher Scientific; Phospho, phosphorylated.

| Antibody | Supplier | Cat. No. | Dilution |
|---|---|---|---|
| RelA | CST | 4764 | 1:1000 |
| Phospho RelA serine 536 | CST | 3031S | 1:1000 |
| RelB | CST | 4954 | 1:1000 |
| c-Rel | CST | 67489 | 1:1000 |
| p100/p52 | CST | 3017 | 1:1000 |
| p105/p50 | 05-361 | Millipore | 1:1000 |
| CYLD | CST | 12797 | 1:1000 |
| Phospho IκBα serines 32/36 | CST | 9246 | 1:1000 |
| EDAR | Protein Tech | 18032-1-AP | 1:1000 |
| GAPDH (mouse) | CST | 97166 | 1:1000 |
| GAPDH (rabbit) | CST | 2118 | 1:1000 |
| **Secondary antibodies** | | | |
| Goat anti-rabbit HRP linked | CST | 7074 | 1:5000 |
| Horse anti-mouse HRP linked | CST | 7076 | 1:5000 |
| IRDye® 680RD donkey anti-mouse | LI-COR | 925-68072 | 1:5000 |
| Alexa Fluor® 790 donkey anti-rabbit | Thermo | A11374 | 1:5000 |

**Immunofluorescence**

Tissue sections from snap frozen skin tumors were fixed in ice cold methanol for 10 min, washed in PBS for 3 x 10 min, 30% sucrose in PBS w/v for 30 min and then blocked in 0.5% BSA, 0.3% Triton X-100 w/v in PBS for 1 h at room temperature. Immunofluorescent labelling with primary antibodies against CYLD, CK14, CK17, CK18, β-catenin and Ki67 was performed overnight at 4°C. The following day, sections were washed with 0.2% BSA, 0.1% Triton X-100 w/v in PBS for 3 x 10 min. DAPI (4′,6-diamidino-2-phenylindole) nuclear stain (0.1 µg/ml) and secondary fluorophore conjugated antibodies were applied for 1 h at room temperature in the dark. After washing 3 x 10 minutes with 0.2% BSA, 0.1% Triton X-100 w/v in PBS, coverslips were mounted with ProLong Gold Antifade (Thermofisher Scientific) and fluorescence images captured by structured illumination microscopy using the Zeiss Axioimager Z2, with Apotome 2.

**Antibodies used for immunofluorescence.** CST, Cell Signaling Technology; BDT, BD Transduction; Jackson, Jackson ImmunoResearch.

| Antibody | Supplier | Cat. No. | Dilution |
|---|---|---|---|
| **Primary antibodies** | | | |
| CK14 | Abcam | Ab7800 | 1:800 |

| | | | |
|---|---|---|---|
| CK17 | Abcam | Ab19067 | 1:100 |
| CK18 | Abcam | Ab668 | 1:200 |
| β-catenin | BDT | 610153 | 1:50 |
| Ki67 | CST | 9449 | 1:400 |
| **Secondary antibodies** | | | |
| Alexa Fluor® 488 goat anti-rabbit | Jackson | 111-545-144 | 1:200 |
| Alexa Fluor® 594 goat anti-mouse | Jackson | 115-585-146 | 1:400 |
| Alexa Fluor® 594 donkey anti-guinea pig | Jackson | 706-585-148 | 1:400 |

**Fluorescence activated cell sorting**

Tissue was cut into 1 $mm^3$ pieces and digested by incubation in 1.5 mg/ml collagenase (Sigma C1889) for 2 h at 37°C with gentle agitation by pipetting every 20 min. DNase I (Sigma 10104159001) was added after one hour at a final concentration of 1 mg/ml. After resuspension in flow buffer (5mM EDTA, 2% FBS w/v in PBS) and passing through a 40μm filter, cells were centrifuged at 900 x g for 5 min and the supernatant aspirated. Red blood cells were removed by resuspending the pellet in red blood cell lysis buffer (Abcam ab204733) and incubating at room temperature for 10 min. Remaining cells were centrifuged at 900 x g for 5 min, resuspended in 2 ml flow buffer and counted. 500 μl was taken from each sample for RNA-seq and kept on ice. The remaining cell suspension was centrifuged at 900 x g for 5 min and the flow buffer aspirated leaving 50 – 100 μl covering the pellet. Cells were resuspended and stained with CD45 antibody conjugated to fluorescein isothiocyanate (FITC) (Thermo Fisher 11-9459-42) at 5μl/$10^5$ - $10^8$ cells in the dark at 4°C for 30 min. During this incubation, the input fractions were centrifuged at 20,000 x g for 3 min, resuspended in 350 μl ice cold RLT buffer from Qiagen's RNeasy Micro Kit (74004) with added 1% 2-Mercaptoethanol and frozen on dry ice. CD45 stained cells were topped up with cold flow buffer to 5 ml and centrifuged at 900 x g for 5 min, resuspended in flow buffer at a maximum of 1 x $10^7$/ml and transferred to a FACS tube. DAPI was added at a final concentration of 3 μM immediately prior to analysis by flow cytometer. Cells were sorted with the BD FACSAria™ II flow cytometer into 200μl FBS. Sorted cells were centrifuged at 4°C for 3 min at 20,000 x g, resuspended in 350μl RLT buffer with 1% 2-Mercaptoethanol and frozen on dry ice. Samples were stored at -80°C in RLT buffer for downstream RNA extraction with the RNeasy Micro Kit (Qiagen 74004) following the manufacturer's protocol.

**Supplementary Table Legends**

**Supplementary Table 1.** Patient samples used in this study.

**Supplementary Table 2.** Candidate non-canonical NF-κB target genes differentially expressed between CD45- CCS tumor and normal cells.

**Supplementary Table 1** - Samples used in this study.

| Patient No | CYLD Genotype | Sex | Age at biopsy | Tissue source | Processing | Assay | Assay sample no |
|---|---|---|---|---|---|---|---|
| 1 | WT | M | 79 | Normal skin | Input | RNAseq | 1 |
| 1 | WT | M | 79 | Normal skin | CD45 positive | RNAseq | 2 |
| 1 | WT | M | 79 | Normal skin | CD45 negative | RNAseq | 3 |
| 2 | WT | M | 77 | Normal skin | Input | RNAseq | 4 |
| 2 | WT | M | 77 | Normal skin | CD45 positive | RNAseq | 5 |
| 2 | WT | M | 77 | Normal skin | CD45 negative | RNAseq | 6 |
| 3 | c.2460delC | F | 77 | Cylindroma | Input | RNAseq | 7 |
| 3 | c.2460delC | F | 77 | Cylindroma | CD45 positive | RNAseq | 8 |
| 3 | c.2460delC | F | 77 | Cylindroma | CD45 negative | RNAseq | 9 |
| 4 | c.2460delC | F | 64 | Cylindroma | Input | RNAseq | 10 |
| 4 | c.2460delC | F | 64 | Cylindroma | CD45 positive | RNAseq | 11 |
| 4 | c.2460delC | F | 64 | Cylindroma | CD45 negative | RNAseq | 12 |
| 5 | c.2460delC | F | 73 | Spiradeno ma | Bulk tissue | RNAseq | 13 |
| 5 | c.2460delC | F | 78 | Cylindrospir adenoma | Bulk tissue | RNAseq | 14 |
| 5 | c.2460delC | F | 78 | Cylindroma | Bulk tissue | RNAseq | 15 |
| 6 | c.2460delC | F | 55 | Cylindroma | Input | RNAseq | 16 |
| 6 | c.2460delC | F | 55 | Cylindroma | CD45 positive | RNAseq | 17 |
| 6 | c.2460delC | F | 55 | Cylindroma | CD45 negative | RNAseq | 18 |
| 7 | WT | F | 67 | Normal skin | Input | RNAseq | 19 |
| 7 | WT | F | 67 | Normal skin | CD45 positive | RNAseq | 20 |
| 7 | WT | F | 67 | Normal skin | CD45 negative | RNAseq | 21 |
| 7 | WT | F | 67 | Normal skin | Input | RNAseq | 22 |
| 7 | WT | F | 67 | Normal skin | CD45 positive | RNAseq | 23 |
| 7 | WT | F | 67 | Normal skin | CD45 negative | RNAseq | 24 |
| 8 | c.2460delC | F | 50 | Cylindroma | Input | RNAseq | 25 |
| 8 | c.2460delC | F | 50 | Cylindroma | CD45 positive | RNAseq | 26 |
| 8 | c.2460delC | F | 50 | Cylindroma | CD45 negative | RNAseq | 27 |
| 8 | c.2460delC | F | 50 | Cylindroma | Input | RNAseq | 28 |
| 8 | c.2460delC | F | 50 | Cylindroma | CD45 positive | RNAseq | 29 |
| 8 | c.2460delC | F | 50 | Cylindroma | CD45 negative | RNAseq | 30 |
| 9 | WT | F | 83 | Normal skin | Bulk tissue | RNAseq | 31 |
| 10 | WT | M | 29 | Normal skin | Bulk tissue | RNAseq | 32 |
| 11 | WT | M | 78 | Normal skin | Bulk tissue | RNAseq | 33 |
| 8 | c.2460delC | F | 50 | Cylindroma | Bulk tissue | RNAseq | 34 |
| 8 | c.2460delC | F | 50 | Cylindroma | Bulk tissue | RNAseq | 35 |
| 12 | c.2469+1A>G | F | 82 | Cylindroma | Bulk tissue | RNAseq | 36 |
| 12 | c.2469+1A>G | F | 82 | Cylindrospir adenoma | Bulk tissue | RNAseq | 37 |
| 3 | c.2460delC | F | 77 | Cylindroma | Bulk tissue | RNAseq | 38 |
| 5 | c.2460delC | F | 73 | Spiradeno ma | Bulk tissue | RNAseq | 39 |

**Supplementary Table 1** - Samples used in this study.

| Patient No | CYLD Genotype | Sex | Age at biopsy | Tissue source | Processing | Assay | Assay sample no |
|---|---|---|---|---|---|---|---|
| 5 | c.2460delC | F | 73 | Cylindroma | Bulk tissue | RNAseq | 40 |
| 11 | WT | M | 78 | Normal skin | Tissue | Immunoblot (RelA, p100/p52) | 1 |
| 11 | WT | M | 78 | Normal skin | Tissue | Immunoblot (RelA, p100/p52) | 2 |
| 13 | c.2460delC | F | 63 | Cylindroma | Tissue | Immunoblot (RelA, p100/p52) | 3 |
| 8 | c.2460delC | F | 47 | Cylindroma | Tissue | Immunoblot (RelA, p100/p52) | 4 |
| 3 | c.2460delC | F | 71 | Cylindroma | Tissue | Immunoblot (RelA, p100/p52) | 5 |
| 13 | c.2460delC | F | 63 | Cylindroma | Tissue | Immunoblot (RelA, p100/p52) | 6 |
| 8 | c.2460delC | F | 47 | Cylindroma | Tissue | Immunoblot (RelA, p100/p52) | 7 |
| 11 | WT | M | 78 | Normal skin | Tissue | Immunoblot (RelB, p105/p50) | 1 |
| 11 | WT | M | 78 | Normal skin | Tissue | Immunoblot (RelB, p105/p50) | 2 |
| 8 | c.2460delC | F | 39 | Cylindroma | Tissue | Immunoblot (RelB, p105/p50) | 3 |
| 8 | c.2460delC | F | 47 | Cylindroma | Tissue | Immunoblot (RelB, p105/p50) | 4 |
| 3 | c.2460delC | F | 71 | Cylindroma | Tissue | Immunoblot (RelB, p105/p50) | 5 |
| 13 | c.2460delC | F | 63 | Cylindroma | Tissue | Immunoblot (RelB, p105/p50) | 6 |
| 8 | c.2460delC | F | 47 | Cylindroma | Tissue | Immunoblot (RelB, p105/p50) | 7 |
| 13 | c.2460delC | F | 61 | Cylindroma | 2D CPCC4 | Immunoblot (CYLD shRNA) | 1 |
| 14 | c.2460delC | F | 44 | Cylindroma | 2D CPCC2 | Immunoblot (CYLD shRNA) | 2 |
| 15 | WT | M | 94 | Normal skin | Tissue | CD45- FACS and proteomics | *C1* |
| 16 | WT | M | 83 | Normal skin | Tissue | CD45- FACS and proteomics | *C2* |
| 17 | WT | M | 69 | Normal skin | Tissue | CD45- FACS and proteomics | *C3* |
| 18 | WT | F | 68 | Normal skin | Tissue | CD45- FACS and proteomics | *C4* |
| 19 | WT | F | 55 | Normal skin | Tissue | CD45- FACS and proteomics | *C5* |
| 20 | WT | F | 67 | Cylindroma | Tissue | CD45-CD200+ enrichment and proteomics | *T1* |
| 21 | c.2460delC | F | 54 | Cylindroma | Tissue | CD45-CD200+ enrichment and proteomics | *T2* |
| 8 | c.2460delC | F | 54 | Cylindroma | Tissue | CD45-CD200+ enrichment and proteomics | *T3* |
| 22 | Sporadic | F | 73 | Cylindroma | Tissue | CD45-CD200+ enrichment and proteomics | *T4* |
| 8 | c.2460delC | F | 54 | Cylindroma | Tissue | CD45-CD200+ enrichment and proteomics | *T5* |
| 23 | WT | M | 56 | Normal skin | Tissue | Immunoblot (CYLD) | 1 |
| 11 | WT | M | 78 | Normal skin | Tissue | Immunoblot (CYLD) | 2 |
| 24 | c.1112C>A | M | 39 | Cylindroma | Tissue | Immunoblot (CYLD) | 3 |
| 25 | 5.5MB deletion | F | 54 | Cylindroma | Tissue | Immunoblot (CYLD) | 4 |
| 25 | 5.5MB deletion | F | 54 | Cylindroma | Tissue | Immunoblot (CYLD) | 5 |
| 8 | c.2460delC | F | 49 | Cylindroma | Tissue | Immunoblot (CYLD) | 6 |
| 26 | c.2469+1G>A | M | 75 | Cylindroma | Tissue | Immunoblot (CYLD) | 7 |
| 27 | WT | M | 71 | Normal skin | Spheroids | Immunoblot (NF-kB) | 1 |
| 28 | WT | M | 83 | Normal skin | Spheroids | Immunoblot (NF-kB) | 2 |
| 29 | WT | M | 68 | Normal skin | Spheroids | Immunoblot (NF-kB) | 3 |
| 8 | c.2460delC | F | 46 | Cylindroma | Spheroids | Immunoblot (NF-kB) | 4 |
| 30 | c.2469+1G>A | M | 51 | Cylindroma | Spheroids | Immunoblot (NF-kB) | 5 |
| 14 | c.2460delC | F | 44 | Cylindroma | Spheroids | Immunoblot (NF-kB) | 6 |
| 29 | WT | M | 68 | Normal skin | 2D NPCC | Immunoblot (CYLD, RelB, Edar) | 1 |
| 31 | WT | M | 71 | Normal skin | 2D NPCC | Immunoblot (CYLD, RelB, Edar) | 2 |
| 3 | c.2460delC | F | 76 | Cylindroma | 2D CPCC | Immunoblot (CYLD, RelB, Edar) | 3 |
| 14 | c.2460delC | F | 44 | Cylindroma | 2D CPCC | Immunoblot (CYLD, RelB, Edar) | 4 |
| 32 | WT | M | 71 | Normal skin | Spheroids | Immunoblot (CYLD, RelB, Edar) | 5 |
| 33 | WT | M | 83 | Normal skin | Spheroids | Immunoblot (CYLD, RelB, Edar) | 6 |
| 8 | c.2460delC | F | 46 | Cylindroma | Spheroids | Immunoblot (CYLD, RelB, Edar) | 7 |
| 30 | c.2469+1G>A | M | 51 | Cylindroma | Spheroids | Immunoblot (CYLD, RelB, Edar) | 8 |

**Supplementary Table 1** - Samples used in this study.

| Patient No | CYLD Genotype | Sex | Age at biopsy | Tissue source | Processing | Assay | Assay sample no |
|---|---|---|---|---|---|---|---|
| 3 | c.2460delC | F | 76 | Cylindroma | Spheroids | Viability (TPCA1) | 1 |
| 5 | c.2460delC | F | 81 | Cylindroma | Spheroids | Viability (TPCA1) | 2 |
| 30 | c.2469+1G>A | M | 51 | Cylindroma | Spheroids | Viability (TPCA1) | 3 |
| 5 | c.2460delC | F | 81 | Cylindroma | Spheroids | Viability (BMS) | 1 |
| 30 | c.2469+1G>A | M | 51 | Cylindroma | Spheroids | Viability (BMS) | 2 |
| 30 | c.2469+1G>A | M | 51 | Cylindroma | Spheroids | Viability (BMS) | 3 |
| 14 | c.2460delC | F | 44 | Cylindroma | Spheroids | Viability (SU1644) | 1 |
| 3 | c.2460delC | F | 76 | Cylindroma | Spheroids | Viability (SU1644) | 2 |
| 30 | c.2469+1G>A | M | 51 | Cylindroma | Spheroids | Viability (SU1644) | 3 |
| 30 | c.2469+1G>A | M | 51 | Cylindroma | 2D CPCC2 | Immunoblot (CYLD +inhibitors) | 1 |
| 3 | c.2460delC | F | 76 | Cylindroma | 2D CPCC3 | Immunoblot (CYLD +inhibitors) | 2 |
| 14 | c.2460delC | F | 44 | Cylindroma | 2D CPCC4 | Immunoblot (CYLD +inhibitors) | 3 |
| 29 | WT | M | 68 | Normal skin | 2D NPCC1 | Immunoblot (CYLD +inhibitors) | 4 |
| 29 | WT | M | 68 | Normal skin | 2D NPCC2 | Immunoblot (CYLD +inhibitors) | 5 |
| 31 | WT | M | 71 | Normal skin | 2D NPCC3 | Immunoblot (CYLD +inhibitors) | 6 |
| 34 | WT | F | 61 | Normal skin | Spheroids | Immunoblot (p100/p52 +SU1644) | 1 |
| 35 | WT | M | 83 | Normal skin | Spheroids | Immunoblot (p100/p52 +SU1644) | 2 |
| 30 | c.2469+1G>A | M | 51 | Cylindroma | Spheroids | Immunoblot (p100/p52 +SU1644) | 3 |
| 13 | c.2460delC | F | 61 | Cylindroma | Spheroids | Immunoblot (p100/p52 +SU1644) | 4 |
| 8 | c.2460delC | F | 53 | Cylindroma | CD45 negative | scRNAseq | 1 |
| 8 | c.2460delC | F | 53 | Cylindroma | CD45 negative | scRNAseq | 2 |
| 8 | c.2460delC | F | 53 | Cylindroma | CD45 negative | scRNAseq | 3 |

**Supplementary Table 2**. Candidate non-canonical NF-κB target genes differentially expressed between CD45$^-$ normal skin and CCS tumour cells.

| No. | $log_2FC$ | Gene | Description |
|---|---|---|---|
| 1 | 8.2 | *EDAR* | Ectodysplasin A receptor |
| 2 | 8.0 | *SOX14* | SRY-box 14 |
| 3 | 5.4 | *KREMEN2* | Kringle containing transmembrane protein 2 |
| 4 | 5.2 | *CD164L2* | CD164 molecule like 2 |
| 5 | 5.2 | *PLA2G4D* | Phospholipase A2 group IVD |
| 6 | 5.0 | *LCP1* | Lymphocyte cytosolic protein 1 |
| 7 | 5.0 | *SOX21* | SRY-box 21 |
| 8 | 4.1 | *COL22A1* | Collagen type XXII alpha 1 chain |
| 9 | 4.0 | *FAM71E1* | Family with sequence similarity 71 member E1 |
| 10 | 3.9 | *RHOV* | Ras homolog family member V |
| 11 | 3.8 | *FAM83F* | Family with sequence similarity 83 member F |
| 12 | 3.5 | *KY* | Kyphoscoliosis peptidase |
| 13 | 3.5 | *FAAH2* | Fatty acid amide hydrolase 2 |
| 14 | 3.4 | *FFAR2* | Free fatty acid receptor 2 |
| 15 | 3.2 | *TSTD1* | Thiosulfate sulfurtransferase like domain containing 1 |
| 16 | 3.2 | *DDR1* | Discoidin domain receptor tyrosine kinase 1 |
| 17 | 3.2 | *DUSP2* | Dual specificity phosphatase 2 |
| 18 | 3.1 | *ANO9* | Anoctamin 9 |
| 19 | 3.1 | *ANK3* | Ankyrin 3 |
| 20 | 2.9 | *KCNB2* | Potassium voltage-gated channel subfamily B member 2 |
| 21 | 2.9 | *VAX2* | Ventral anterior homeobox 2 |
| 22 | 2.9 | *ELF3* | E74 like ETS transcription factor 3 |
| 23 | 2.6 | *LHFPL4* | LHFPL tetraspan subfamily member 4 |
| 24 | 2.5 | *THSD7B* | Thrombospondin type 1 domain containing 7B |
| 25 | 2.5 | *NMBR* | Neuromedin B receptor |
| 26 | 2.5 | *STAP2* | Signal transducing adaptor family member 2 |
| 27 | 2.5 | *HEPACAM* | Hepatic and glial cell adhesion molecule |
| 28 | 2.4 | *ZNF385C* | Zinc finger protein 385C |
| 29 | 2.4 | *DAPP1* | Dual adaptor of phosphotyrosine and 3-phosphoinositides 1 |
| 30 | 2.4 | *TRAF1* | TNF receptor associated factor 1 |
| 31 | 2.4 | *SAMD5* | Sterile alpha motif domain containing 5 |
| 32 | 2.4 | *ARHGAP4* | Rho GTPase activating protein 4 |
| 33 | 2.3 | *BSN* | Bassoon presynaptic cytomatrix protein |
| 34 | 2.3 | *SGPP2* | Sphingosine-1-phosphate phosphatase 2 |
| 35 | 2.2 | *GOLT1A* | Golgi transport 1A |
| 36 | 2.2 | *AGBL2* | ATP/GTP binding protein like 2 |
| 37 | 2.1 | *GNA15* | G protein subunit alpha 15 |
| 38 | 2.1 | *HPCA* | Hippocalcin |
| 39 | 2.0 | *RNF227* | Ring finger protein 227 |
| 40 | -2.0 | *CLGN* | Calmegin |
| 41 | -2.1 | *HTRA1* | HtrA serine peptidase 1 |

| No. | $log_2FC$ | Gene | Description |
|---|---|---|---|
| 42 | -2.2 | *CLDN5* | Claudin 5 |
| 43 | -2.3 | *PTGDS* | Prostaglandin D2 synthase |
| 44 | -2.3 | *RRAD* | Ras related glycolysis inhibitor and calcium channel regulator |
| 45 | -2.4 | *ACKR1* | Atypical chemokine receptor 1 (Duffy blood group) |
| 46 | -2.5 | *RSPO1* | R-spondin 1 |
| 47 | -2.9 | *RIMS2* | Regulating synaptic membrane exocytosis 2 |
| 48 | -3.6 | *TOR4A* | Torsin family 4 member A |
| 49 | -4.0 | *ITIH1* | Inter-alpha-trypsin inhibitor heavy chain 1 |

Applying a threshold of fold change ($log_2FC$) 2 and adjusted *P*-value < 0.05, 49 differentially expressed genes contain a p52:RelB binding site +/-1kb from the transcription start site. Positive FC indicates expression is higher in CCS tumour CD45- cells relative to CD45- cells from normal skin.

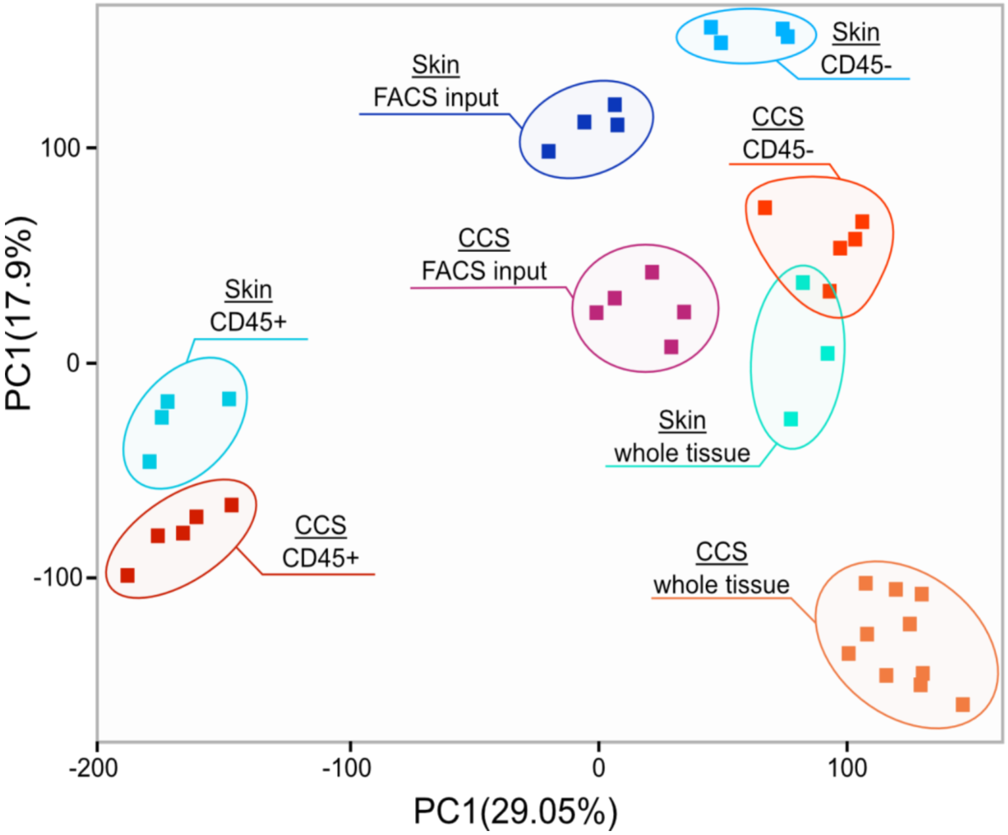


**Supplementary Figure 1. Bulk RNA sequencing of CCS tumours and CD45$^{+}$ and CD45$^{-}$ cell fractions.** Principal component analysis of transcriptomic data from FACS sorted CCS tumours reveals distinct clusters of CD45$^{+/-}$ CCS and CD45$^{+/-}$ normal cells.

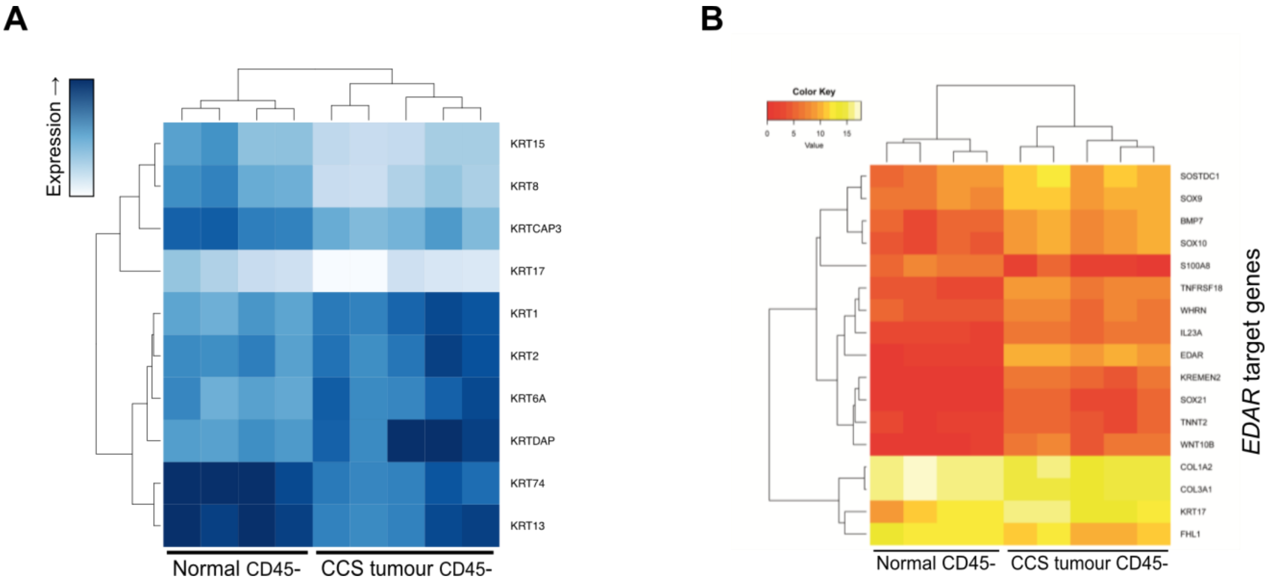


**Supplementary Figure 2. CD45- CCS tumour cells have a different cytokeratin profile to CD45-normal skin cells. (A)** CD45- cells isolated from CCS tumors and normal skin by FACS differentially express 10 protein coding keratin genes (log2FC>2, adjusted P-value <0.05). Dark blue indicates low expression, light blue indicates high expression. The gene encoding keratin 74 (KRT74), a hair follicle specific keratin, was ranked first by positive fold-change. (B) CD45- cells isolated from CCS tumours and normal skin by FACS differentially express 16 potential EDA/EDAR signalling pathway target genes (FC2, adjusted P-value < 0.05). Heatmap shows hierarchical clustering by Euclidean distance. Red indicates low expression, light-yellow indicates high expression.

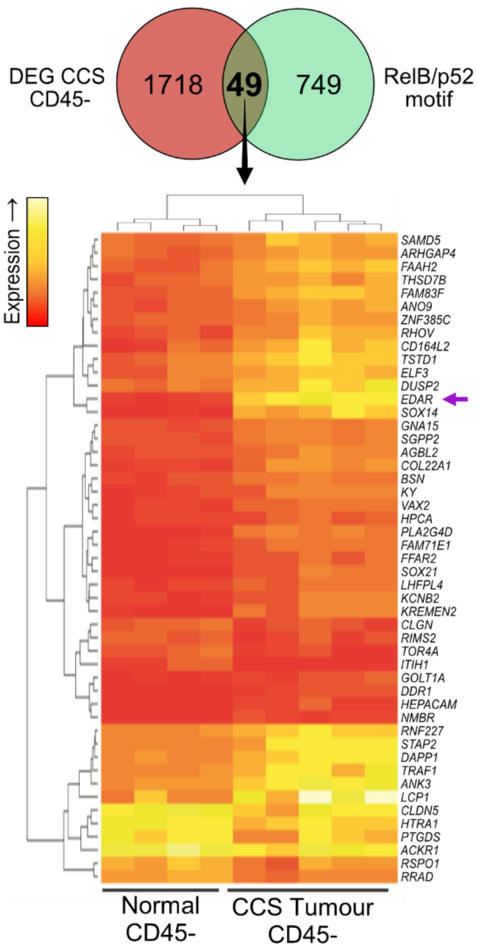


**Supplemental Figure 3. An EDAR cluster is upregulated in CD45⁻ CCS tumor cells** 49 genes are differentially expressed between FACS sorted CD45⁻ CCS tumor and CD45⁻ normal skin cells containing the p52/RelB motif GGGGNTTTCC. Hierarchical clustering by Euclidean distance of gene expression produces clusters within a heatmap. Red indicates low expression, light-yellow indicates high expression. *EDAR* had the greatest positive fold-change from 1718 DEGs (FC>2, adjusted P-value < 0.05) (purple arrow).

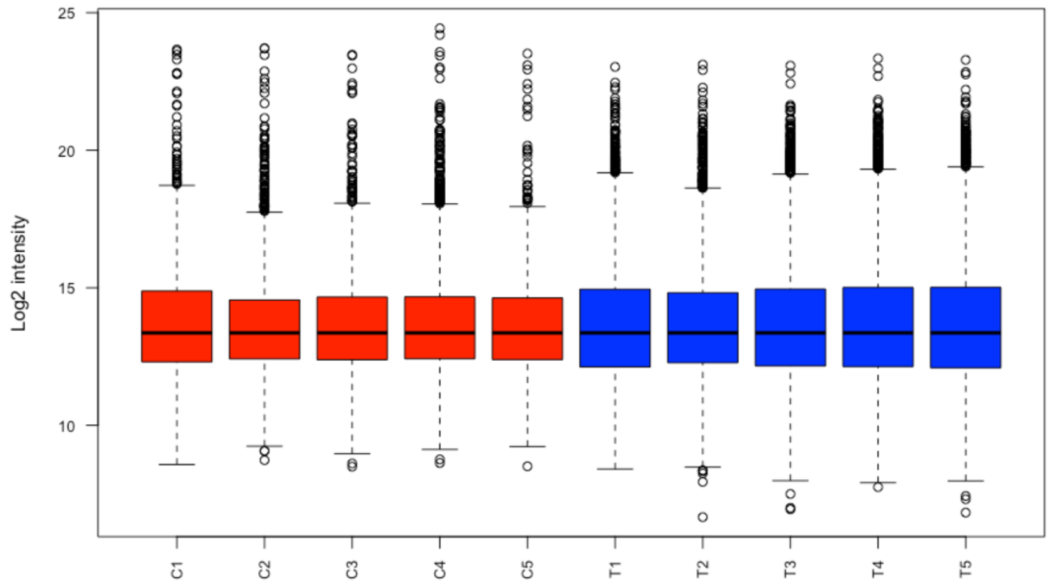


**Supplementary Figure 4. Normalised log$^2$ intensities of proteins detected in CD45$^-$ epidermis and CD45$^-$/CD200$^+$ CCS tumour cells.** Median normalised log$^2$ intensities of CD45$^-$ epidermis (C1-C5) and CD45$^-$/CD200$^+$ CCS tumour cells (T1-T5).

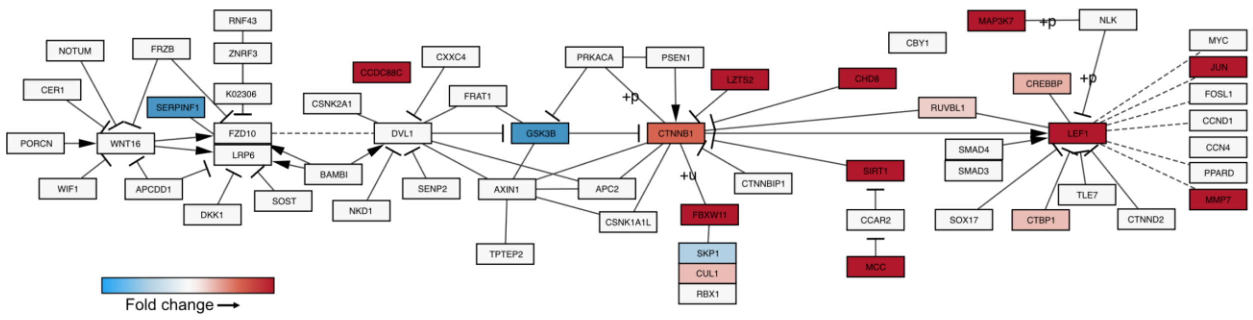


**Supplementary Figure 5. KEGG pathway of Wnt-Beta catenin signalling with differential protein expression between CCS CD45-CD200+ and normal skin CD45- cells overlaid.** Fold change is indicated as increasing (red) or decreasing (blue). Non-coloured proteins were not detected.

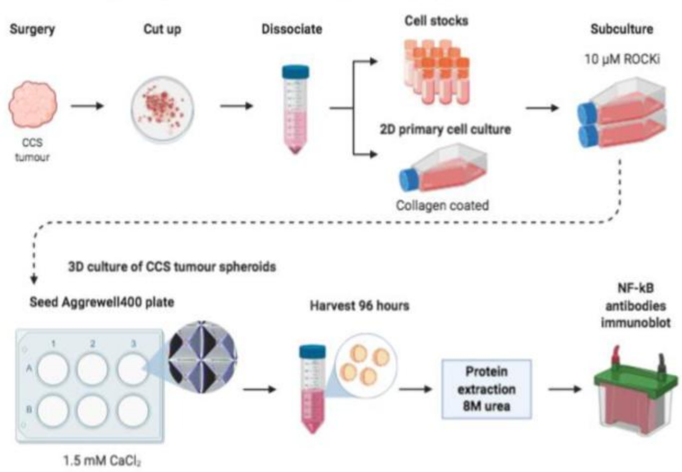


**Supplementary Figure 6. 3D culture of primary CCS tumour keratinocytes.** CCS tumours were dissociated in 2.5% trypsin and 1 mg/ml collagenase at 37°C. 2D cultured cells were seeded on collagen coated flasks and subcultured in keratinocyte-serum free media (KSFM) supplemented with 10 μM Y-27632 (ROCKi). Two confluent T75 flasks were pooled to seed an Aggrewell400 plate containing 7000 microwells per well. Within 24h, one 3D spheroid aggregate formed per microwell. Spheroids were harvested at 96h and protein extracted by sonication in 8M urea lysis buffer for immunoblot analysis of NF-κB signalling.

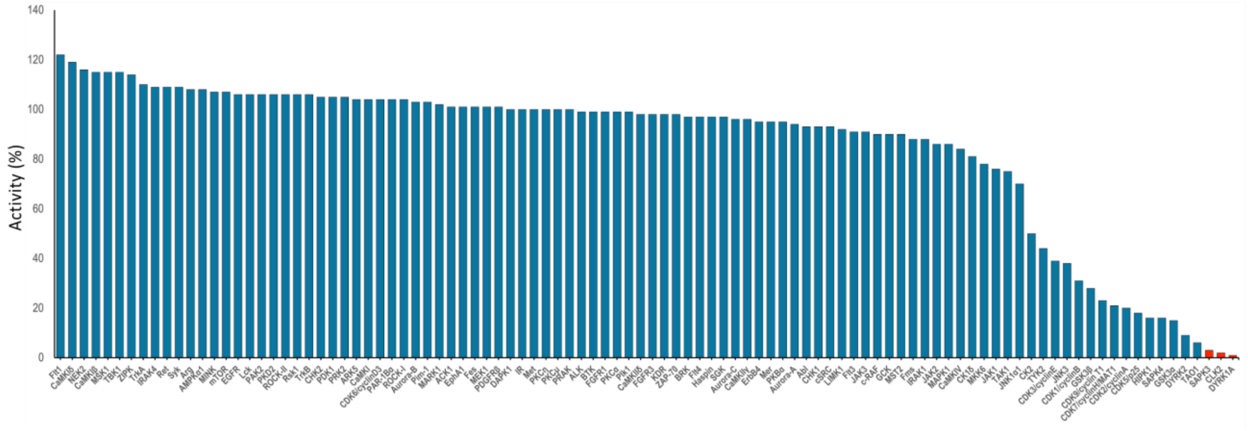


**Supplementary Figure 7. SU1644 demonstrates selectivity for IKKalpha over other kinases.** Kinome profile for SU1644: % residual kinase activity @ 1 μM (at ATP Km)

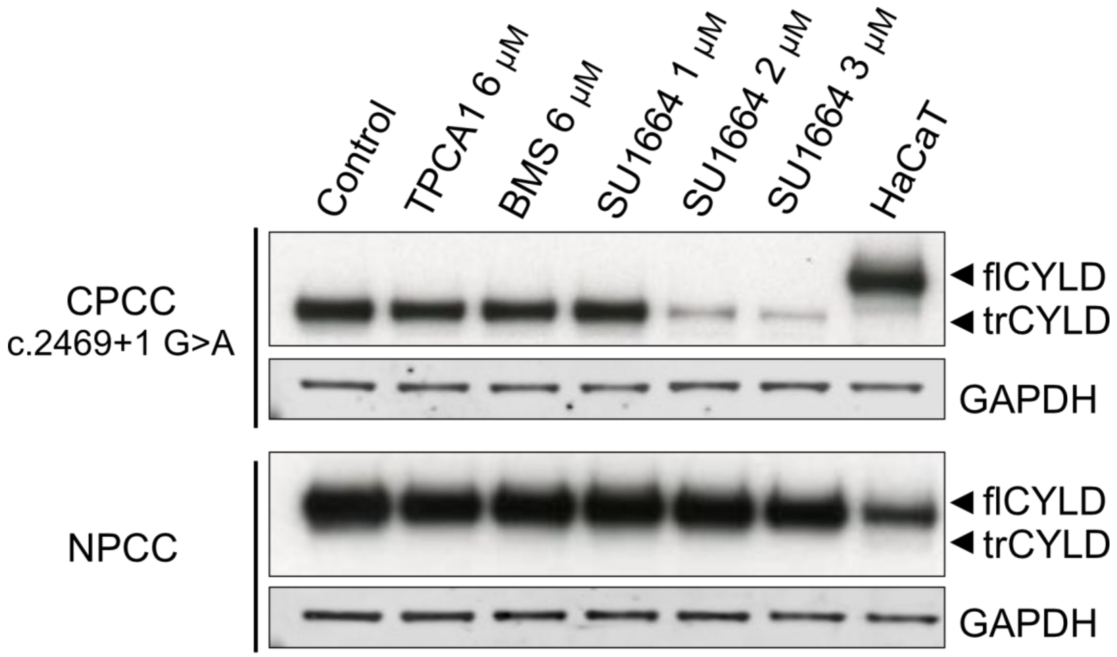


**Supplementary Figure 8. Truncated CYLD levels are reduced in a splice site pathogenic variant of *CYLD* after SU1644 treatment.** Levels of trCYLD (truncated CYLD) and flCYLD (full length CYLD) in CCS primary cell culture (CPCC) and normal primary cell culture (NPCC) detected by immunoblotting after 24 h treatment with IKK inhibitors. HaCaT lysate is included as a positive control for flCYLD. BMS; BMS345541.
