## Supplementary material for "Targeting non-canonical NF-κB signalling in CYLD cutaneous syndrome by selective inhibition of IκB kinase alpha": Unedited Western blot images are available in a separate file

Full unedited gel for

### Figure 3A

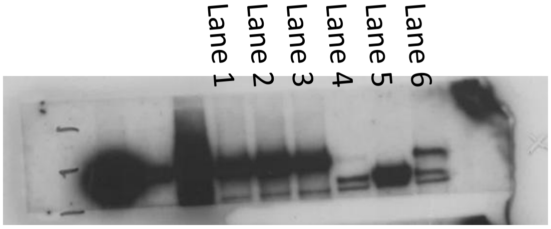


CYLD

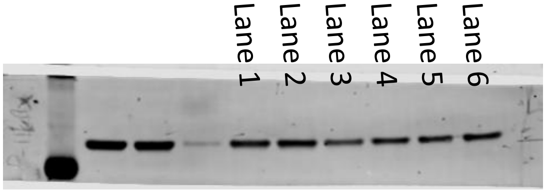


GAPDH

Full unedited gel for

### Figure 3B-C

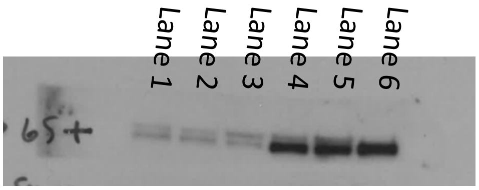


**RelB**

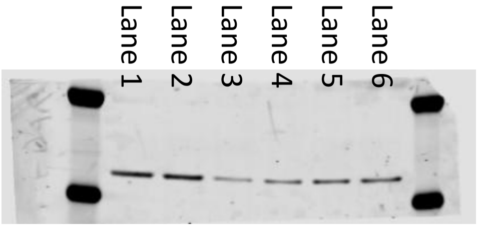


**GAPDH – 1 (RelB)**

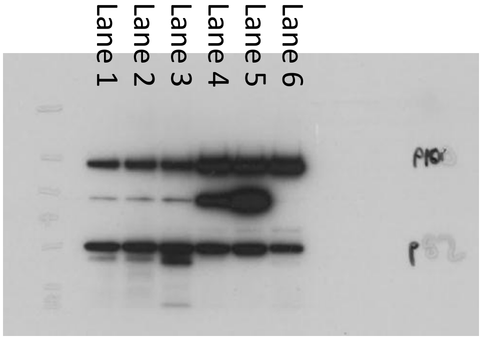


**p100/p52**

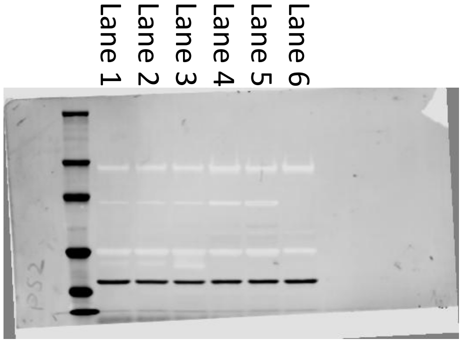


**GAPDH – 2 (p100/p52)**

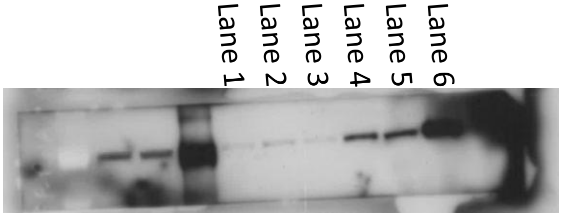


***p*-RelA**

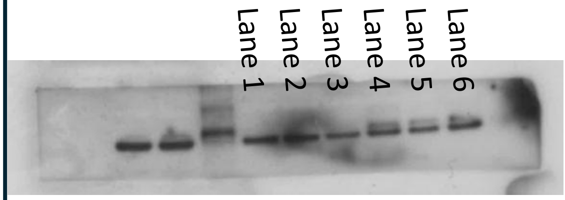


***p*-IκBα**

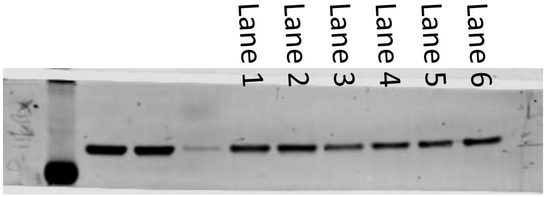


**GAPDH – 3 (*p*-RELA, *p*-IκBα )**

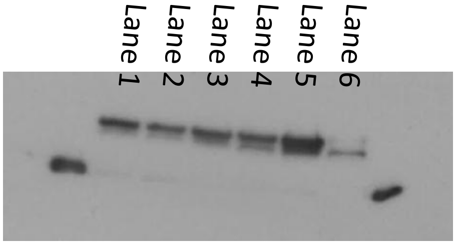


**RelA**

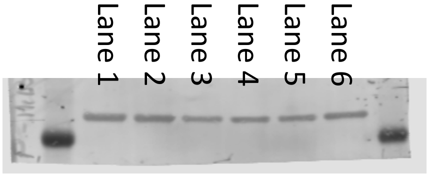


**GAPDH – 4 (RelA)**

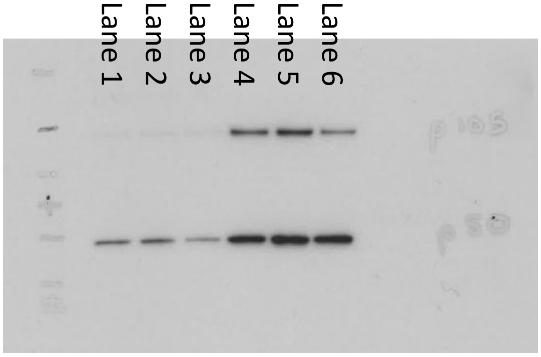


**p105/p50**

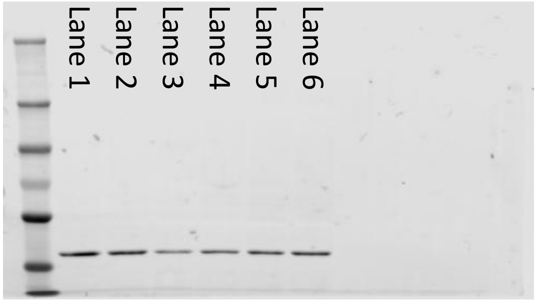


**GAPDH – 5 (p105/p50)**

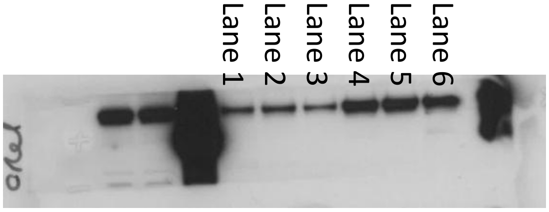


**c-Rel**

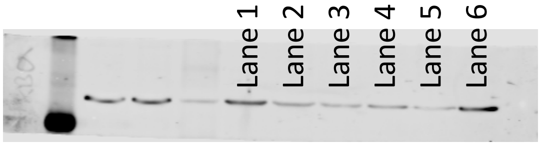


**GAPDH – 6 (c-Rel)**

Full unedited gel for

### Figure 3D

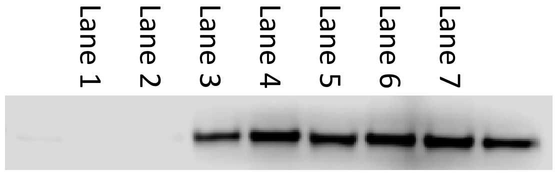


**p100**

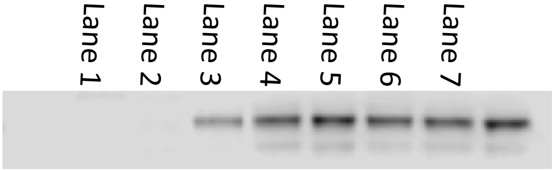


**p52**

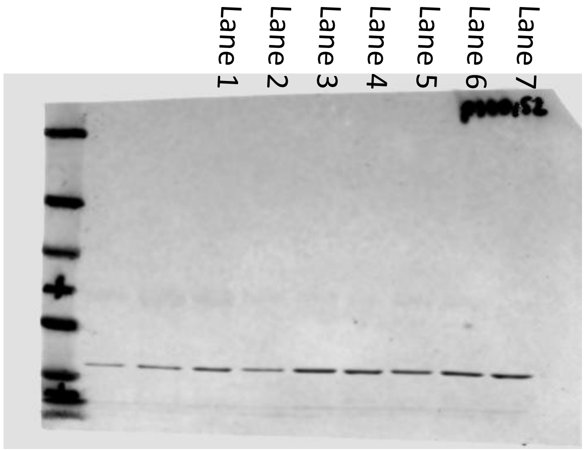


**GAPDH – 1 (p100/p52)**

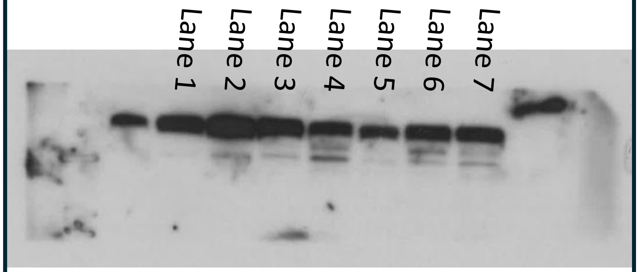


**RelB**

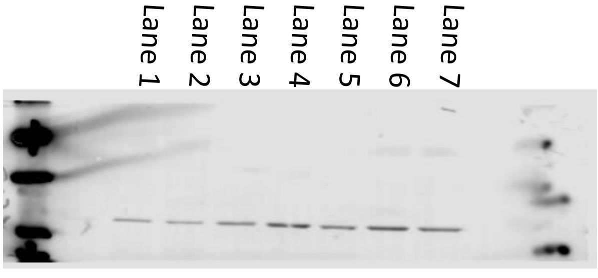


**GAPDH – 2 (RelB)**

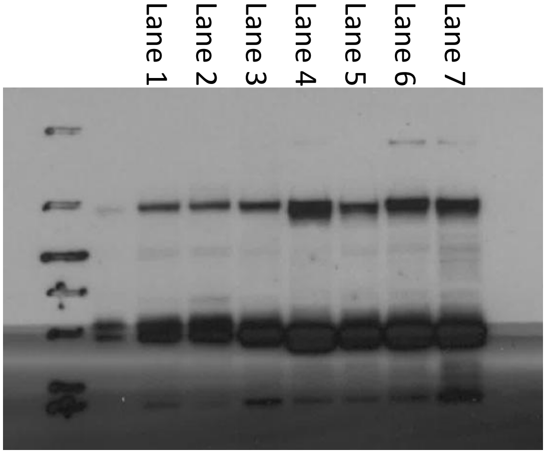


**p105**

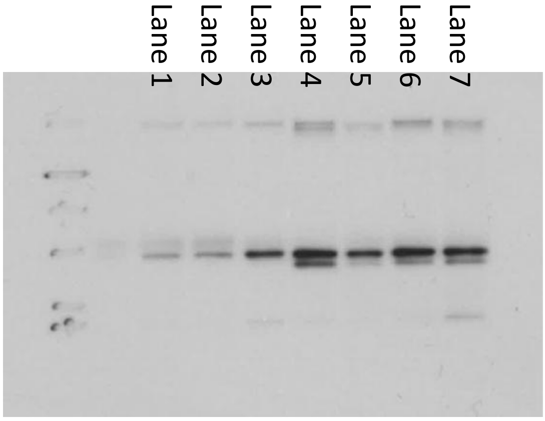


**p50**

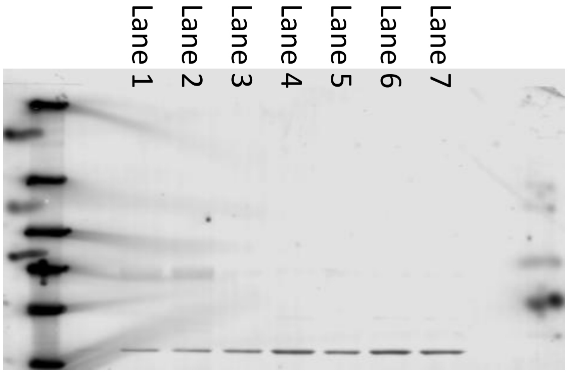


**GAPDH – 3 (p105/p50)**

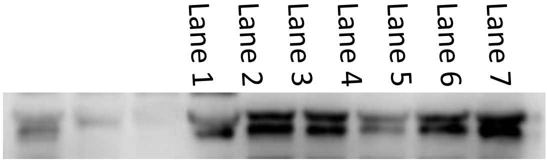


**RelA**

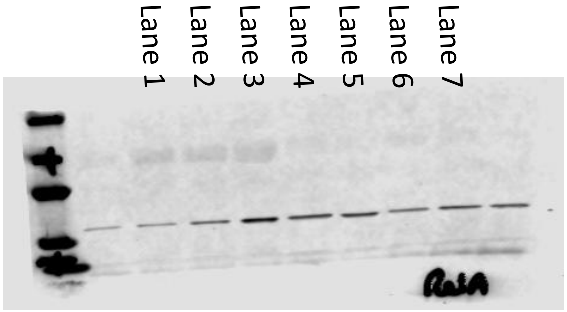


**GAPDH – 4 (RelA)**

Full unedited gel for

### Figure 4D

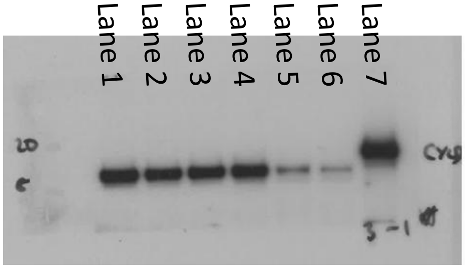


**CYLD – CPCC1**

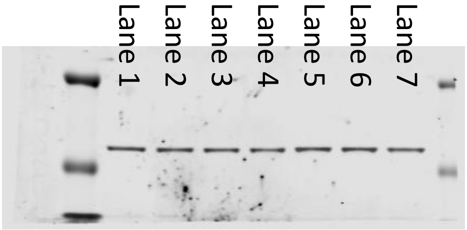


**GAPDH – CPCC1**

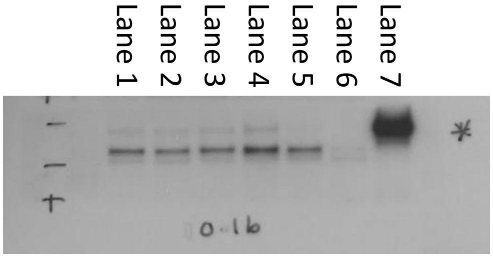


**CYLD – CPCC2**

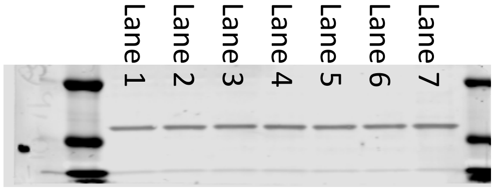


**GAPDH – CPCC2**

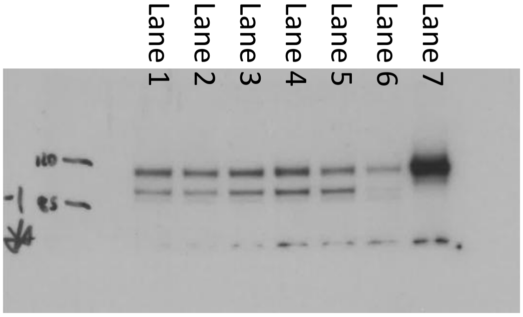


**CYLD – CPCC3**

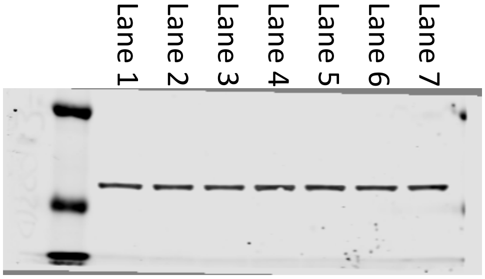


**GAPDH – CPCC3**

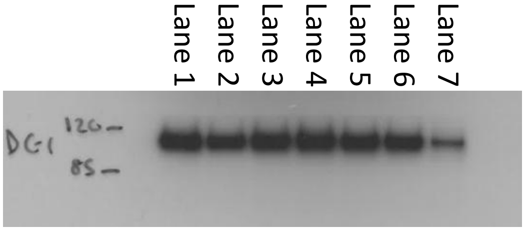


**CYLD – NPCC1**

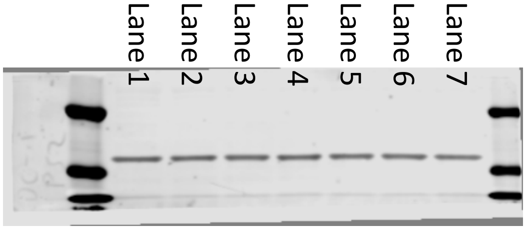


**GAPDH – NPCC1**

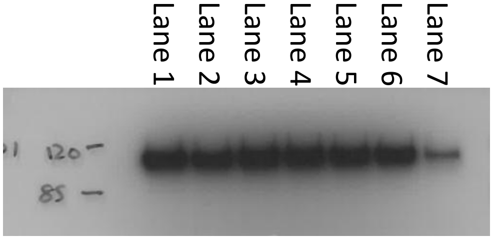


**CYLD – NPCC2**

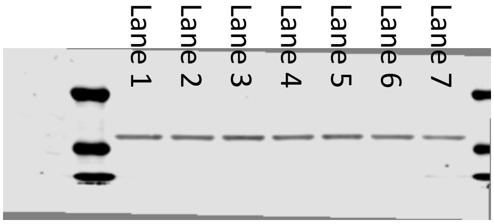


**GAPDH – NPCC2**

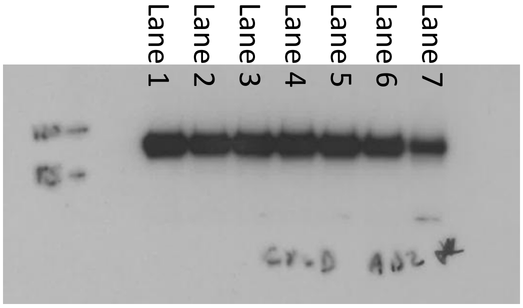


**CYLD – NPCC3**

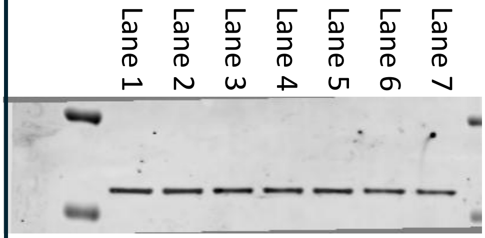


**GAPDH – NPCC3**

Full unedited gel for

### Figure 4E

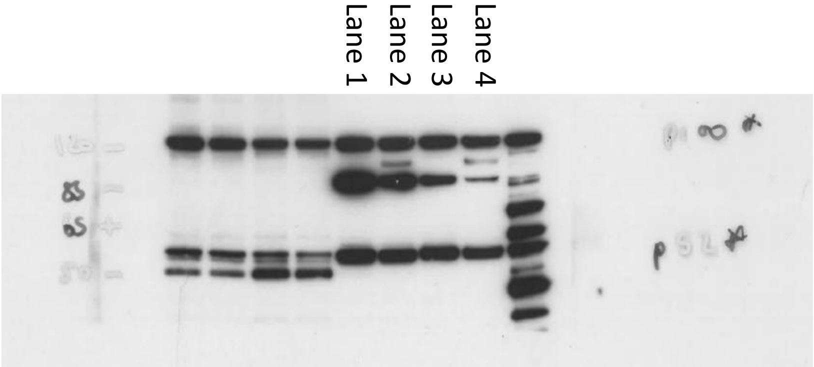


**p100/p52 (60 s)**

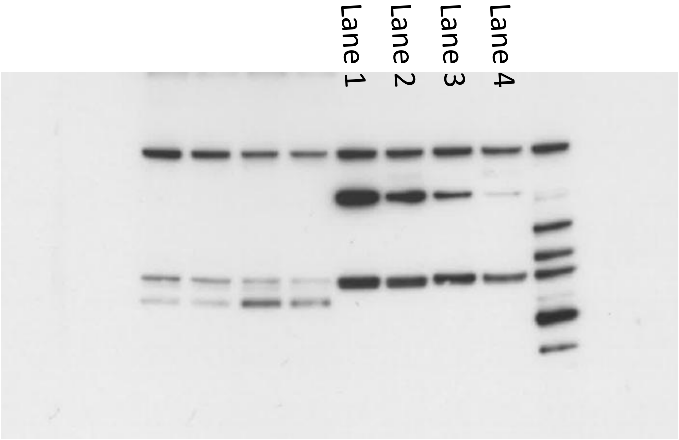


**p100/p52 (10 s)**

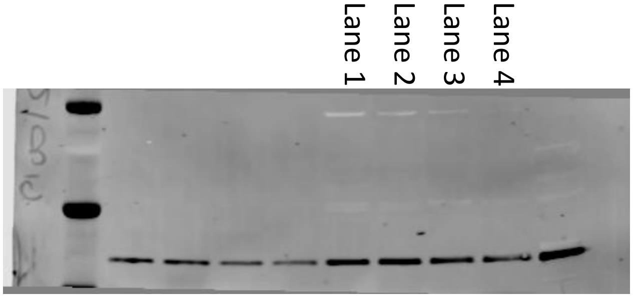


**GAPDH**
